# MnM: a machine learning approach to detect replication states and genomic subpopulations for single-cell DNA replication timing disentanglement

**DOI:** 10.1101/2023.12.26.573369

**Authors:** Joseph M. Josephides, Chun-Long Chen

## Abstract

We introduce MnM, an efficient tool for characterising single-cell DNA replication states and revealing genomic subpopulations in heterogeneous samples, notably cancers. MnM uses single-cell copy-number data to accurately perform missing-value imputation, classify cell replication states and detect genomic heterogeneity, which allows to separate somatic copy-number alterations from copy-number variations due to DNA replication. By applying our machine learning methods, our research unveils critical insights into chromosomal aberrations and showcases ubiquitous aneuploidy in tumorigenesis. MnM democratises single-cell subpopulation detection which, in hand, enables the extraction of single-cell DNA replication timing (scRT) profiles from genomically-heterogenous subpopulations detected by DNA content and issued from single samples. By analysing over 119,000 human single cells from cultured cell lines, patient tumours as well as patient-derived xenograft samples, the copy-number and replication timing profiles issued in this study lead to the first multi-sample subpopulation-disentangled scRT atlas and act as data contribution for further cancer research. Our results highlight the necessity of studying *in vivo* samples to comprehensively grasp the complexities of DNA replication, given that cell lines, while convenient, lack dynamic environmental factors. This tool offers to advance our understanding of cancer initiation and progression, facilitating further research in the interface of genomic instability and replication stress.

**GRAPHICAL ABSTRACT:** 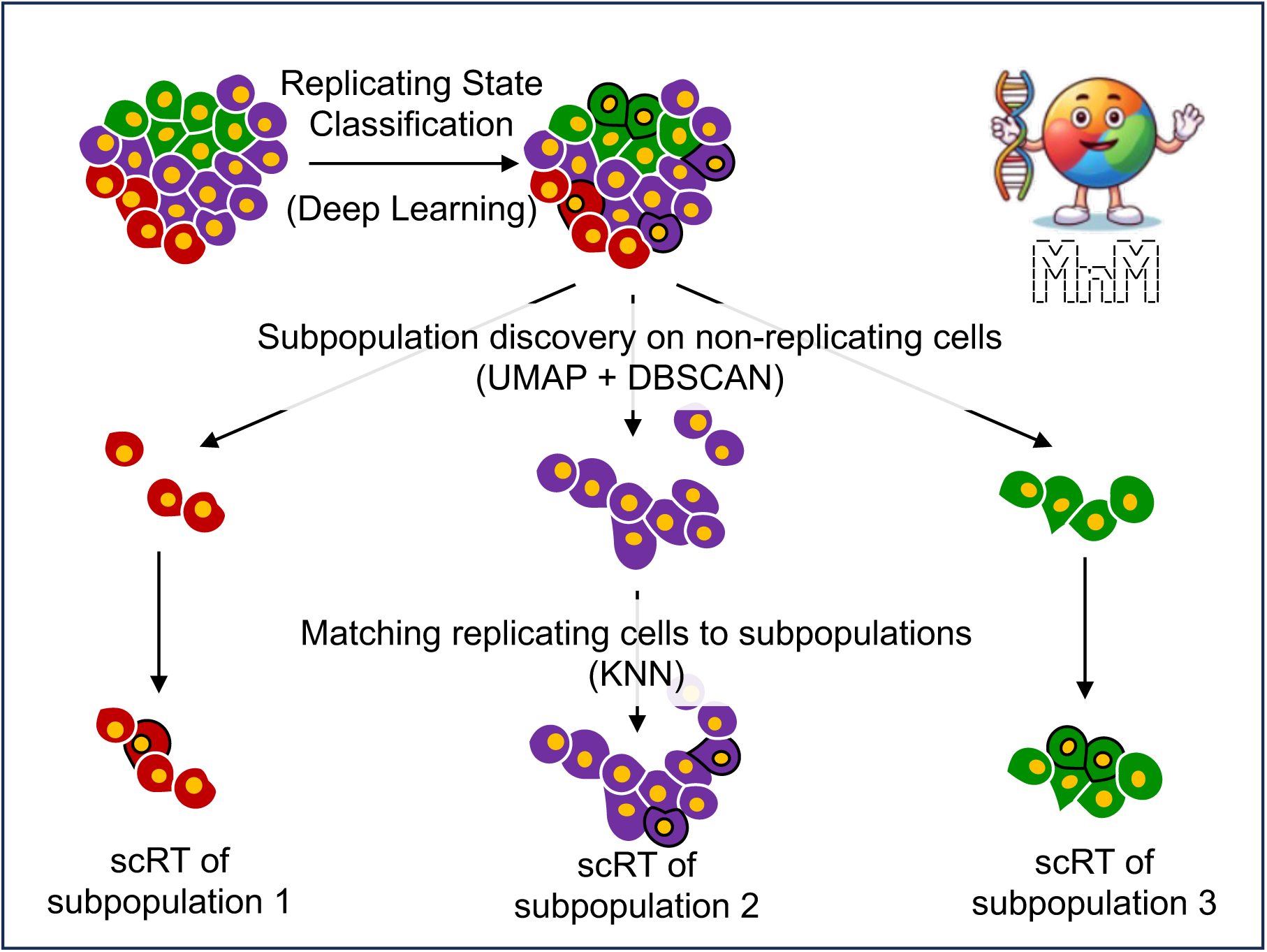

## INTRODUCTION

DNA replication is a fundamental biological process in which, under normal circumstances, a cell creates an identical copy of its genome to ensure accurate transmission of genetic information to the daughter cells during the synthesis (S)-phase of the cell cycle. Despite the robustness and molecular orchestration of this process, errors can occur leading to inexact, over-or under-replication. Thus, the generation of genomic alternations including DNA copy number alterations (CNAs) (1–4), point mutations (5–7), and other structural variants is common (8–13). CNAs refer to abnormal fluctuations in the number of copies of specific genomic regions within a cell’s genome and can arise due to various cellular processes, including DNA replication errors, lack of replication factors, chromosomal instability, and DNA damage (14). Dysregulation of DNA replication is associated with various human diseases, including cancer (15, 16). In fact, studies have shown that replication stress, which occurs when cells encounter obstacles that prevent the successful execution of the replication program, can cause genomic instability and promote tumorigenesis, making it a hallmark of cancer (14, 17). Therefore, the study of CNAs and the DNA replication process is particularly important in cancer research, as they can impact gene expression and function, potentially leading to tumour initiation, progression, therapy resistance and serving as biomarkers for cancer diagnosis, prognosis, and treatment response (18, 19). However, there is a lack of methods that are adapted to study DNA replication specifically in cancerous cell populations.

The traditional way to detect CNAs from next generation sequencing (NGS) data was by measuring the number of DNA copies per genomic region from bulk sequencing. Bulk whole-genome sequencing was, and remains, a common approach to detect CNAs in cancer samples. However, this technique presents limitations in terms of genomic heterogeneity detection, an important aspect of evolving tumour populations (11, 20). With bulk sequencing, the genetic material from multiple cells is mixed and sequenced, which only provides an average image of the genomic alterations across the tumour cells and neglects subclones with distinct genomic profiles which may be critical for understanding both the origin and evolution of CNAs in cancer. Therefore, bulk sequencing, by overlooking the degree of genomic heterogeneity in tumours, restricts the identification of genomic changes that are critical for studying cancer progression, drug resistance and understanding therapeutic bottlenecks.

Recently, the advent of single-cell technologies has revolutionised cancer research. Single-cell genomics, in particular, has enabled the study of cancer heterogeneity and the identification of rare cell populations that may be responsible for tumour initiation and therapy resistance. This technology has been used to characterize CNAs and other genomic alterations in individual cells, revealing the extent of intra-tumoral heterogeneity and clonal evolution in cancer (20–28). While the discovery of genomic subpopulations, groups of cells that have distinct CNA signatures in comparison to other cells originating from the same sample, has been enabled, it is overlooked when studying DNA replication.

A key metric in studying DNA replication is Replication Timing (RT), a critical aspect of S-phase that refers to the order in which different regions of the genome are copied during the cell cycle. RT is highly regulated and correlates with other cellular processes, including gene expression (29), DNA methylation (30), chromatin structure (31) as well as the 3D organisation of chromosomes (32). With single-cell whole genome sequencing (scWGS) upgrading RT studies, more detailed investigations in mammalian replication dynamics at the genome-wide level have emerged (30, 32–36). Until recently, single-cell RT (scRT) investigations were limited by the number of cells that could be studied in each sample due to technical restraints. Some studies that have overcome these hurdles (30, 35, 36) have confirmed that cell-to-cell heterogeneity in RT can be found in a single sample. We have previously shown that it is possible to distinguish subpopulations from CNAs and extract distinct replication patterns using scWGS data issued from a single heterogenous cell line sample (35). However, this process was not automated and consequently, RT is not routinely disentangled in tumours.

In order to democratise RT studies in complex cancers, we developed a new tool – Mix ‘n’ Match (MnM) – an automated machine learning-based tool that exploits CNAs of single cells from mixed cell populations to match cells together based on similarity, reconstructing subpopulations, while also discovering replication states *in silico*. By grouping cells based on their cell cycle phase and genomic makeup, we were able to observe distinguished RT trajectories for subpopulations issued from cell lines and tumour samples. We rigorously trained a ready-to-use supervised machine learning model to identify which cells are replicating and applied unsupervised learning to identify subpopulations. By analysing 119,991 human single-cells, we provide the largest single source to date of heterogeneity-resolved scRT profiles.

## MATERIALS AND METHODS

### MnM: Mix ‘n’ Match single-cells

MnM consists of three main steps: passing single-cell copy-number data through (i) k-Nearest Neighbors (KNN) imputation to complete missing copy-number values, (ii) a supervised replication state classifier (separating replicating from non-replicating cells) and then (ii) through an unsupervised subpopulation detector (see below for details). The program can load a bed file containing DNA copy-numbers per genomic region (obtained with scWGS data) with cell indices or, alternatively, a matrix with genomic regions as headers (chr:star-end) and individual cell identifiers as row indices (Fig. 1A). Missing copy-numbers are filled-in with KNN imputation (Fig. 1B), and the replication states of the cells in the data are predicted from a pre-trained and ready-to-use deep learning model (Fig. 1C,D). Copy-numbers of non-replicating cells are then subjected to a low-dimensional space transformation to discover underlying subpopulations through unsupervised clustering (Fig. 1E). Finally, the replicating cells are matched with their corresponding non-replicating populations on a reduced dimension landscape (Fig. 1F) allowing further analysis, such as scRT extraction or focusing on CNAs of the non-replicating cells of the different subpopulations. A representation of MnM’s main steps (Fig. 1) illustrates that the combination of deep learning, Uniform Manifold Approximation and Projection (UMAP), Density-based spatial clustering of applications with noise (DBSCAN) and KNN algorithms allows uncovering replication states and subpopulations from single-cell whole-genome copy-number calling data (detailed in the following sections).

**Figure 1.**
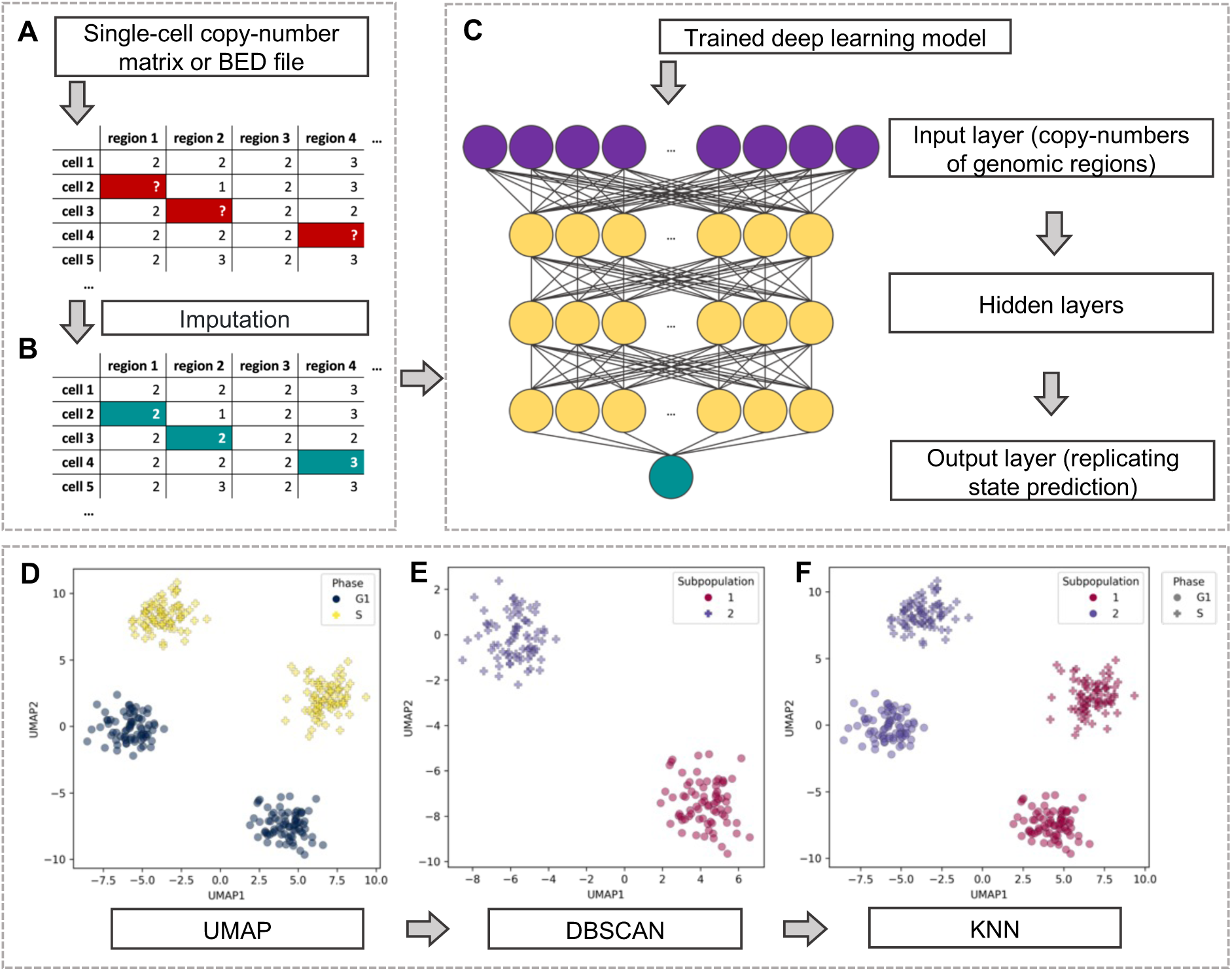
Machine learning techniques used throughout MnM. **(A-B)** Copy-number imputation with k-Nearest Neighbors (KNN). Single-cell copy-number data is used as an input in either a matrix or BED file format with missing copy-number values (A) which are filled in with KNN imputation (B). **(C)** Deep learning for a single-cell replication state classifier. The trained deep learning model consisting of 3 hidden and 1 output layer is then loaded and used to distinguish replication states of the single cells. **(D-F)** Subpopulation discovery in 3 steps. Dimensionality reduction is performed with UMAP on non-replicating cells under 2 dimensions to provide representative lower dimensions of the copy-number data (D). DBSCAN clusters the data based on the UMAP coordinates (E) which allows replicating cells to be matched to the corresponding non-replicating subpopulations with KNN under 10-dimension UMAP coordinates (F).

### scWGS demultiplexing

The BAM files of hg19-aligned GM12878 B lymphocyte cell-line from ref. (36) were obtained, sorted by read name with samtools (37) sort (v. 1.16.1; option -n) and converted to fastq files with samtools fastq (options -T CB --barcode-tag CB) to have barcodes transcribed in the fastq headers from the BAM headers to be aligned to the hg38 reference genome as indicated below. These files were demultiplexed with demultiplex (38) demux (v 1.2.2; options -m 0 -- format x). Other cells obtained from the 10X Chromium Single Cell CNV solution from ref. (26, 30, 36) (Supplementary Table 1-3) were acquired as fastq files and demultiplexed using demultiplex demux (options -r -e 16) with barcodes extracted from the first 16 bp of forward reads. All extracted barcodes were filtered based on the 10X barcode whitelist.

### Barcode validation

10X Single-cells were considered for analysis if they originated from valid barcodes which were identified as follows. Data was prepared by counting the number of lines of each demultiplexed fastq file and then divided by 4 to reflect the number of total reads per single-cell. The resulting list containing the number of reads per barcode was then used to make a distinction between corrupted or low-read (invalid) barcodes from qualitative (valid) ones through a custom R (39) (v4.0.4) script. Barcodes containing less than 30,000 reads were considered to not be qualitative due to the very low number of reads, were systematically removed to eradicate any noise in the initial peak and with the goal of only keeping a mixture of two distributions (valid and invalid barcodes). The em command from the cutoff R library (v0.1.0) was used to identify the cut-off point of 2 log-normal distributions of the read counts from the Expectation-Maximisation (EM) algorithm for each demultiplexed file.

EM was comprised of two stages, the expectation (E-step) and maximisation (M-step) steps, which occurred after initialisation of the μ and σ parameters (see below) for the 2 log-normal distributions (D1 and D2). The probability density function (PDF) used during the E-step, which represented the probability of observing a particular read count per barcode (continuous random variable) given the following parameters, of the log-normal distribution can be described as:

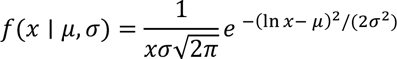

where:

- 𝑥 represents the read count of the barcode.
- μ represents the mean (also called location parameter).
- σ represents the standard deviation (also called the scale parameter).

Using this, 𝛾_i_ which represents the probability that barcode read count 𝑖 belongs to the valid distribution is calculated as:

where:

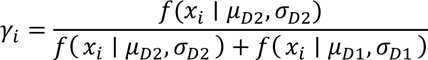

- 𝑓(𝑥_i_ ∣ 𝜇_𝐷2_, 𝜎_𝐷2_) is the PDF of the log-normal distribution with parameters 𝜇_𝐷2_, 𝜎_𝐷2_ evaluated at the read count 𝑥_i_ for the valid distribution (D2).
- 𝑓(𝑥_i_ ∣ 𝜇_𝐷1_, 𝜎_𝐷1_) is PDF of the log-normal distribution with parameters 𝜇_𝐷1_, 𝜎_𝐷1_ evaluated at the read count 𝑥_i_ for the invalid distribution (D1).

This E-step computed the expected value of any missing data points and calculated the probabilities of the missing or overlapping data given the estimates of μ and σ. The following M-step then updated the parameters of the log-normal distributions using the estimated probabilities as follows:

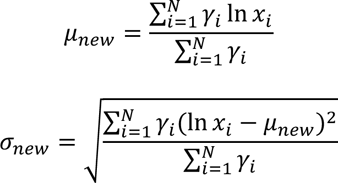

The E- and M-steps were repeated iteratively until the estimated probabilities 𝛾_i_converged (when the parameters and probabilities stopped changing between iterations). The exact cut- off value between D1 and D2 was obtained with the cutoff command from the same package, having D1, the lower read count distribution, belonging to the Type-I error. Only the barcodes having a number of reads superior or equal to the EM cut-off value were considered to be valid and those with a lower number of reads were discarded. Histograms containing representations of the read counts and the cut-off values were systematically generated for visual inspection and validation. The valid barcodes were retained with their respective reads as fastq files corresponding to single-cells used for mapping and further analysis.

### Read mapping

Single-cell WGS data of MCF-7 breast cancer (35), JEFF B lymphocyte (35) and HeLa S3 cervical cancer cells (35), along with hTERT-RPE1 retinal pigment epithelial cells (33) were aligned to the UCSC human reference genome hg38 as previously reported (35) using the Kronos FastqToBam module. Other single-cell fastq files had their reads trimmed and filtered by quality score with Trim Galore (40) (v0.6.4; options –fastqc, –gzip, --paired when paired-end data were used or omitted otherwise, and --clip_R1 16 except for the GM12878 cells that were originally aligned to hg19) based on Cutadapt (41) (v3.7) and FastQC (42, 43) (v0.11.9) and mapped onto the UCSC hg38 reference genome with BWA mem (44) (v0.7.17; option -M). Mate coordinates were corrected using samtools fixmate (option -O bam) whenever the data were issued from paired-end sequencing. All BAM files were then sorted by coordinates with samtools sort (-O bam) before read duplicates were removed with Picard (45) MarkDuplicates (v2.26.11; options ASSUME_SORT_ORDER=coordinate, METRICS_FILE) via java (v19; options - Xmx16g -jar). MultiQC (46) (v1.10.1) was used to visually inspect single-cell quality.

### Copy-number matrix organisation

Copy-numbers from the resulting single-cell BAM files were estimated with the Kronos scRT Binning and CNV commands in either 20 or 25 kb windows (see code for details). Problematic genomic regions were masked with the v2 hg38 ENCODE blacklist (47). The resulting copy-number BED files were regrouped by sample and used as an input for further analyses and visual representations with MnM with the random seed set to 18671107.

Genomic regions from all MnM input files were rearranged in 100 kb non-overlapping genomic windows (as a median of the copy-numbers from the input file which overlapped the 100 kb window by at least 50%) delimited by the chromosome sizes of the hg38 reference genome provided by bedtools (48, 49), and in 25 kb and 500 kb replication state classifier models. MnM then automatically processed the data by temporarily removing windows containing no data and any remaining sporadic missing values were filled in with the integrated sklearn KNN imputation algorithm (50) (options n_neighbors=5, weights=’distance’). The nearest neighbours were defined as the 5 closest cells based on the Euclidean distance of the genome-wide copy-numbers (distances calculated in pairs for genomic regions that neither of the 2 cells were missing). A weighted average of copy-numbers from the region of the closest neighbours was used as the imputation value. The imputation method can be described as:

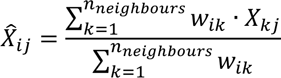

where:

- 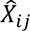 represents the imputed value for the copy-number of the region 𝑗 in cell 𝑖.
- 𝑋_𝑘𝑗_ denotes the value of region 𝑗 in the 𝑘-th neighbour.
- 𝑛*_neighbours_* is the number of nearest neighbours considered for imputation. Here 𝑛 =5.
- 𝑤_i𝑘_ represents the weight assigned to the 𝑘-th neighbour for cell 𝑖 based on their Euclidean distance.

This imputation method was also used for the imputation of 5-55% in intervals of 5% of single-cell copy-number values that were randomly selected and removed after the elimination of any windows containing missing values of an S-phase-enriched population of MCF-7 cells (35). A random imputation method where each missing copy-number value was substituted by a randomly selected non-missing value from the matrix, along with a median imputation method where the median of each genomic region was imputed, were implemented for comparison to the KNN imputation method under the same random seed (see code for details). Accuracy was calculated as the percentage of identity of the imputed values compared to the original values. Similarity was calculated as the percentage of values that differed less than ±1 copy number for KNN imputation compared to the original value. Invariance was calculated matrix-wide as the percentage of unchanged copy-numbers after imputation.

### Replication state classifier

To organise the data for the replication state classifier, cells phases were either extracted with Kronos for HeLa, MCF-7 and JEFF cells (35), solely from the FACS metadata for hTERT-RPE1 cells (33) or from the intersection of common replication states from the FACS metadata and Kronos for HCT-116 colon cancer and sorted GM12878 cells (30, 36) (Supplementary Table 2). The resulting single-cell copy-number matrices were concatenated. Any partially or completely missing regions (i.e. any genomic region containing at least one missing copy-number value) were removed while only autosomal data were retained. Eighty percent of the cells were used as training data and the remaining 20% were used as testing data. To prepare the replication state classifier to be able to distinguish noisy copy-number non-replicating profiles (e.g. from low-quality cells or technical noise) from those of replicating cells, data augmentation was performed as follows to reduce overfitting. Half of the training cells were randomly selected and copied. For each of these copied cells, noise was induced by altering the copy-numbers by ±1 between 5-75% of the genomic regions, which were selected from a uniform distribution.

The replication state classifier was built on a Sequential architecture, a feed-forward neural network. The model was designed with the Keras (51) python library (v2.13.1) to facilitate the construction of a linear stack of neural network layers, each connected to the subsequent one. As an input, the single-cell copy-number matrix of the training dataset containing the 6 cell types along with the simulated data was used. The sequence of layers aimed at hierarchical feature extraction and predictive modelling consisting of three hidden layers with 64, 32 and 16 units, respectively. These layers facilitated the extraction of increasingly complex and abstract representations of the input copy-number profiles. The model terminated in an output node having a single unit, using a sigmoid activation. This configuration was suited for binary classification tasks, enabling the model to produce a probability estimation in a [0,1] range. Upon construction, the model was compiled with a binary cross-entropy loss function to optimise the network’s performance concerning binary classification. An ‘adam’ optimiser, known to be efficient and adaptive on learning rates, was used to optimise the parameters throughout training. In order to avoid overfitting, an early stopping mechanism was implemented on an epoch-based patience of 15 iterations.

With the completion of training, the resulting neural network model along with the list of genomic windows comprised in the matrix were saved. The model was then integrated and automatically loaded with MnM to predict the single-cell binary replication states (Replicating/S-Phase, Non-Replicating) of the scWGS data obtained from cell line, patient tumour and PDX samples. In the case where any regions required by the model were not present, MnM compensated for these missing values by using linear interpolation from both directions. Compensating for these missing values ensured the continuity and integrity of replication state predictions.

### Subpopulation discovery

Commencing with non-replicating cells, the number of variables were reduced from the number of autosomal regions to 2 dimensions with UMAP (52). The DBSCAN algorithm (53, 54) was then used to detect the number of groups on this reduced dataset with min_samples=10% of the total number of cells. The epsilon (𝜖) parameter was calculated as:

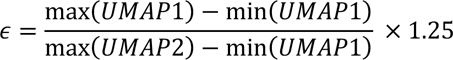

where UMAP1 and UMAP2 correspond to UMAP’s first and second output parameters, respectively. Epsilon was always restricted between 1.25 and 2 while a minimum of at least 10 cells was required to form a subpopulation. UMAP was repeated with 6 randomly generated seeds and the most frequent number of subpopulations, as defined with DBSCAN, was retained. Subpopulations discovered with DBSCAN were redefined and merged iteratively in a descending similarity order if the median copy numbers per region were 98.5% identical. Copy-numbers of both S-phase and non-replicating cells were reduced to 10 UMAP dimensions (second round of UMAP) and then each S-phase cell was matched to the closest non-replicating group with the sklearn nearest neighbour command (options n_neighbors=50% of cells, metric=’euclidean’). The number of nearest neighbours was required to have a minimal value of 5 cells. Both rounds of UMAP were performed on the single-cell copy-number matrices with the addition of 5 artificial cells stretching from complete haploid to pentaploid profiles for subpopulation calibration.

### DNA replication timing

Kronos scRT was modified to work with R v4.0.5, ignore copy-number confidence during quality-control filtering and produce an extra metadata file containing cell diagnostic details. We used the diagnostic module at a first stage for quality control based on the number of reads per Mb under developer mode (option -d) created for this purpose. The data were filtered and passed through MnM for replication state classification and subpopulation detection. The Kronos scRT WhoIsWho module was used to assign the cell phases from the replication state classifier or FACS data accordingly (see code for details) followed by the diagnostic module, which was used a second time to correct the early and late S-phase copy-numbers (option -C). For each subpopulation and biological replicate, the copy-number data were split into different files with a custom python (v3.9.11) code. Kronos scRT was then used to calculate the replication timing profiles through the RT module in 200 kb windows. The resulting scRT binary values were used to produce scRT trajectories with the DRed module under the random seed ‘18671107’ for reproducibility. Permutation tests on the trajectories were made using a custom python code under 1,000 permutations. The observed test statistic was calculated as the absolute mean of sum of differences in means between subpopulations for both UMAP coordinates:

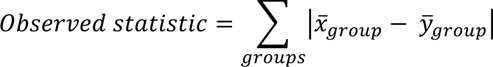

Where *x̄_group_* and *ȳ_group_* are the means of the UMAP1 and UMAP2 coordinates for each subpopulation, respectively.

The permutation test was executed by randomly shuffling the group labels by keeping the UMAP coordinates intact. The test statistic was computed on the shuffled data in the same way as the observed statistic. This allowed to calculate a p-value as follows:

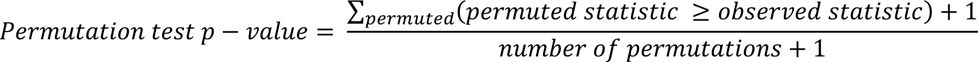

Pseudo-bulk, from scRT, and bulk RT correlations were calculated with the Spearman method and scRT correlation clustering for the scRT atlas was ordered with the Ward.D2 hierarchical clustering method. Bulk RT profiles were lifted over from hg19 to hg38 with the UCSC liftover tool (55) after being converted to bed files with bigwigtobedgraph (56).

## RESULTS

### Highly accurate data completeness for unsynchronised single-cell copy-numbers

Due to technical limitations, single-cell whole-genome sequencing data frequently has a lower read coverage across the genome (i.e. <1X, Supplementary Table 3) in comparison to bulk experiments, leading to sporadic missing copy-number values. To address this issue, we used the KNN imputation technique – a data completion method that takes into consideration the closest cells in terms of genome-wide copy-number profiles (50). To take into account rare CNAs and replication events, we used a weighted copy-number distance for the KNN imputation, which generated an imputed value proportional to the closeness of the genome-wide copy-number profiles between the cells based on Euclidean distances. For each missing value, the existing copy-number for that same region from the closest 5 single cell profiles were used to fill in the missing data (see methods for details).

We proceeded to empirically validate this method by simulating sparce single-cell copy-number matrices. We introduced random voids within the 100 kb window single-cell matrix of replicating and non-replicating MCF-7 cells (n=2,321; 1,288 genomic regions), a cell-line with a large number of CNAs (35), after removing any regions that already contained missing values. We removed between 5% to 55% random values in increments of 5%. We observed that KNN imputation predicted and integrated these missing values with an average accuracy of 83.959%, thereby reconstructing the single-cell copy-number landscape. Remarkably, our findings showed an invariance rate, defined as the total percentage of intact values of the whole matrix, ranging between 99.209% and 90.911% for 5% to 55% of missing values, respectively (Supplementary Fig. 1). These values were significantly higher (paired t-test, p-values<10^-16^) than median (ranging between 97.433% and 71.776%) or random (ranging between 96.11% and 57.27%) imputations, underscoring the robustness of the KNN approach. Furthermore, the imputed values with an absolute difference no larger than 1, in comparison with the original values, ranged between 99.917% and 99.015%, illustrating that the vast majority of the errors introduced through this process were not radically inaccurate, even when more than half of the dataset contained missing values.

To extract the most information possible from our data, we applied the KNN imputation method to all datasets in this study (Supplementary Table 1). Imputed values accounted for merely 0.84%, 0.68% and 1.11% missing values for HeLa, JEFF and MCF-7 cells, respectively, suggesting that the imputed value fidelity for these samples was high (>99%) based on the simulations.

### Deep learning single-cell DNA replication state classifier with high accuracy

Considering that sorting cells by their replication state with fluorescence-activated cell sorting (FACS) can induce errors (57), it would therefore be of interest to further validate replication states by other means. Current computational methods that do so (20, 35, 36) require manually established thresholds or supplementary information such as GC content and intra-cellular variability measurements, which are not always directly accessible. To create a method that can bypass any need of metadata, we amalgamated single-cell copy-numbers issued from datasets with replication states inferred from either FACS (33), Kronos scRT (35), or the intersection of both (30, 36) depending on appropriate extraction methods and harvested the labelled replication states to create a deep learning model based solely on single-cell DNA copy-numbers. A total of 5,250 replicating and 2,273 non-replating cells spanning amongst 6 cell-lines resulted from the amalgamation (Supplementary Table 2). We hypothesised that the various ploidy landscapes of the selected cell lines would make this prediction tool universal and thus adapted to any ploidy state.

We split our dataset in an 80:20 proportion to create training and test datasets. To prepare the model to handle noise from lower quality datasets, the training dataset was augmented with half of its cells replicated and artificially altered to induce random noise of ±1 copy-number sporadically. We trained the model for copy-numbers in 25, 100 and 500 kb bins which resulted in 97.94, 98.54 and 98.14% replication state classification accuracy rates on the test datasets, respectively. To quantify how well our 100 kb model performed in comparison with FACS sorting, we calculated the discordance percentage between our in-silico predictions and the FACS metadata of wild-type (WT; n=713), double-knockout of maintenance DNA methyltransferase DNMT1 and *de novo* DNA methyltransferase DNMT3B (DKO1; n=668) HCT-116 and GM12878 (n=3,180) cells. We observed that FACS misclassifications accounted for 17.67% of WT, 27.70% of DKO1 HCT-116, and 25.72% of GM12878 cells (Fig. 2). These results demonstrate that even when taking into consideration the 1.56% error rate of our model, it generated results with accuracy superior to FACS for cell-phase sorting, in accordance with previous estimations (36).

**Figure 2.**
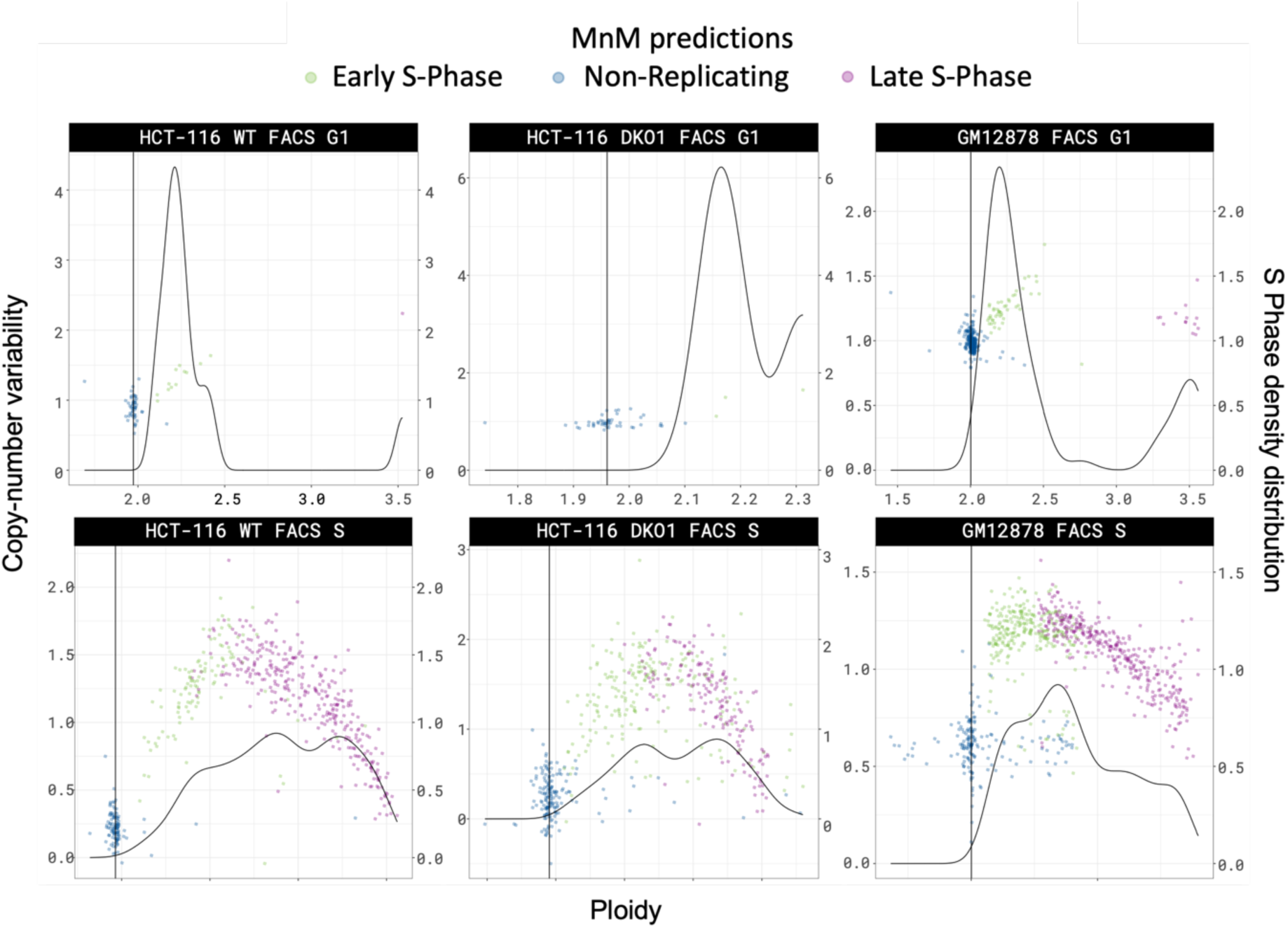
Widespread misclassification of DNA replication states by FACS for HCT-116 WT, DKO1 and GM12979 cell lines. Partial discordance among FACS and the supervised deep learning method developed here on replication states of single-cells. The upper row corresponds to FACS sorting of G1 cells and the lower pane of S cells of HCT-116 wild-type (WT; left), double knock-out (DKO1; centre) and GM12878 (right) cells.

### Unsupervised machine learning automates subpopulation discovery of cancerous cells

Our next goal was to detect copy-number differences between cells, a crucial aspect of cancer emergence and evolution. To do so, we created a 3-step framework to detect genomic subpopulations – groups of cells that have distinct CNA signatures in comparison to other cells originating from the same sample (Fig. 1). The autosomal copy-numbers of non-replicating cells, determined by our replication state classifier, underwent dimensionality reduction to be represented on a two-dimensional pane. The 2D cell coordinates in these new representations would then be used to detect subpopulations with DBSCAN, an unsupervised spatial clustering algorithm. Although UMAP is relatively stable, due to it being a stochastic algorithm (52) that could generate non-representative distances of high-dimensional data, the UMAP/DBSCAN steps were repeated another 6 times under random seeds ranging between 3 and 2^30^. The number of sub-populations was counted for each seed. If the predominate number of clusters was not found in the original iteration, the seed would change to the first encountered of the 6 random seeds that would. Subpopulations were iteratively merged while they presented >98.5% median copy-number identity in a prioritised (decreasing identity) order. Finally, replicating cells would also be included and a second dimensionality reduction step in 10 dimensions allowed the KNN algorithm to match replicating cells to their corresponding non-replicating subpopulations.

To validate this method on genome-wide distinct CNA landscapes, we first mixed JEFF (n=1,455) and HeLa (n=752) copy-number data to be analysed as if they were a single sample, with the expectation that the two cell lines would be correctly distinguished. We first observed that the replicating cells in both cell lines were visually distinguishable in the 2D landscape (Supplementary Fig. 2A-C). After running our 3-step subpopulation detector, we confirmed that, without providing any information on the cell origins, they were matched back into 2 populations corresponding to JEFF and HeLa for both non-replicating (Supplementary Fig. 2d) and replicating (Supplementary Fig. 2C) cells with high accuracy (99.83% and 99.82% for HeLa and JEFF cells, respectively). Furthermore, we exposed the existence of only copy of chromosome X of JEFF (Supplementary Fig. 2D) cells, instead of two which is typical for females, a phenomenon compatible with acquired monosomy X.

In our previously publication (35), we reported the revelation of 2 subpopulations of MCF-7 cells, a breast cancer cell-line known for unstable aneuploidy. We used our current process to automatically detect subpopulations from the MCF-7 sample (n= 2,768) from a single origin. The 2 subpopulations could be distinguished by sub-chromosomal (Fig. 3A) and whole-chromosome (Fig. 3B) copy-number differences and were divergent on UMAP’s reduced dimension pane (Fig. 3E). We then applied this same method to HCT-116 colon carcinoma cells and discovered the existence of 2 subpopulations (Fig 3F), which was previously unreported (30). Contrary to the MCF-7 cells, these subpopulations could only be distinguished on a sub-chromosomal level (Fig 3C-D), suggesting that the observed local CNA changes were due to DNA repair pathways rather that global genome instability. This agrees with the fact that the HCT-116 cell line is known to be defective for the mismatch repair pathway (containing a homozygous mutation of the mismatch repair gene hMLH1 on chromosome 3) and to exhibit microsatellite instability (58, 59).

**Figure 3.**
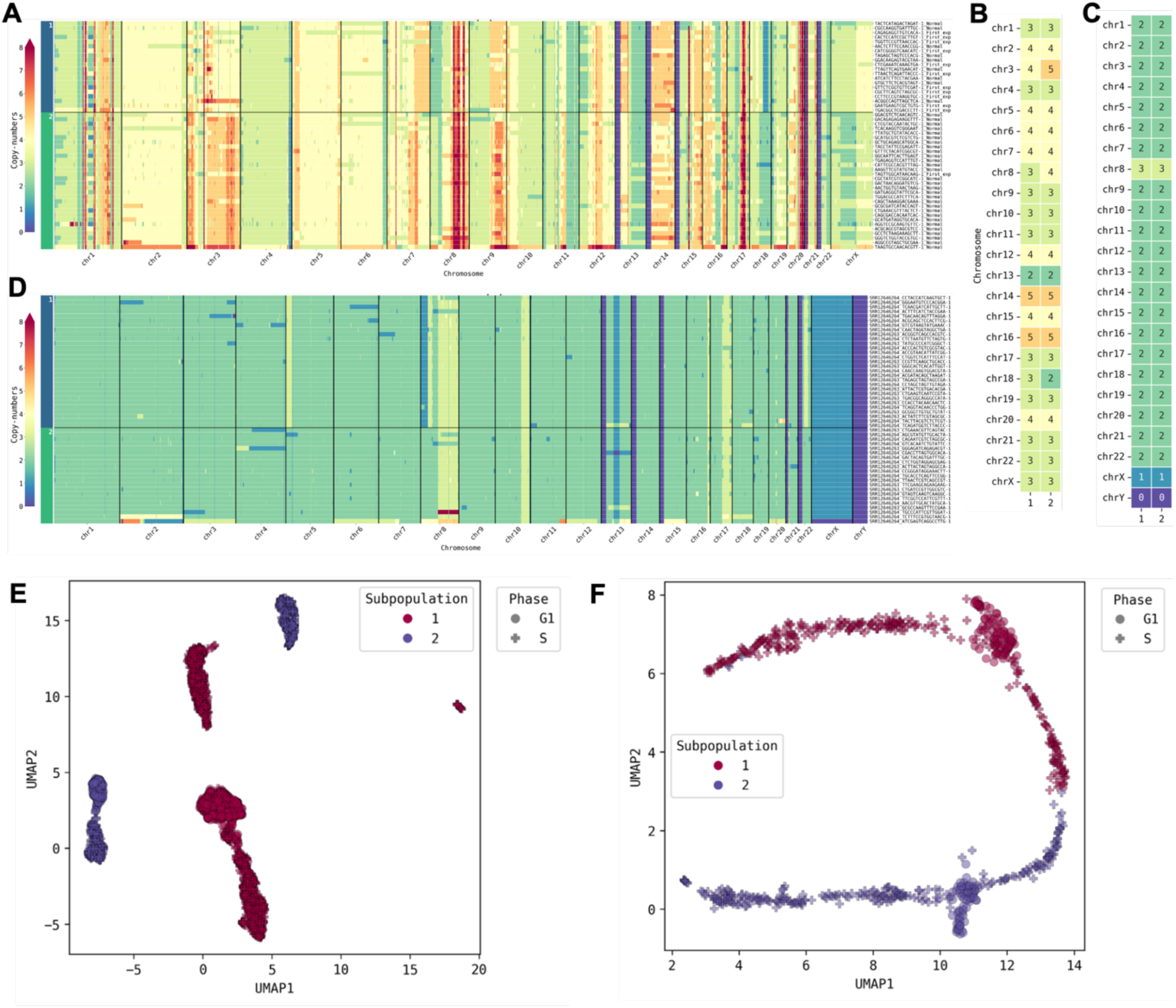
Genomic heterogeneity detected in individual samples of cancer cell lines. **(A-D)** genome-wide copy-numbers (A,D) summarised by their median (B-C) of MCF-7 (A-B) and HCT-116 (C-D) cells. **(E-F)** reduced dimension planes by UMAP of MCF-7 (E) and HCT-116 (F) cells.

To further test whether our subpopulation discovery method can be applied to scWGS data obtained and analysed with different techniques, we reanalysed published copy-number data of 43,106 cells processed in (24), with many originating from (20), which were aligned to hg19 and generated with HMMCopy in 500 kb bins, a reference genome and copy-number estimator which were different to those of the data using for training the deep learning model. Upon visual inspection of single-cell genome-wide copy-number heatmaps (Supplementary Fig. 3), we once again observed that there were copy-number signatures specific to each subpopulation. Thus, we concluded that our approach is robust as it is efficient on different reference genomes and with scWGS data obtained with different techniques (10x scCNV solution, DPL+, etc.) as well as different copy number calling methods (HMMCopy and Kronos scRT).

### DNA replication timing retains high fidelity in cell-lines despite CNAs

We then split the copy number data by subpopulation and provided the detected cell phases to Kronos scRT to obtain the RT profiles. Because the copy-numbers calculated were relative, the first and second parts of S-phase copy-numbers were corrected for MCF-7 (Fig. 4A-B) and HCT-116 (Fig. 4C-D) cells in 200 kb bins. We observed that the S/G1-phase borderline was non-linear on the bin-to-bin variability scale (Fig. 4A-D), signifying that separation of the replication states with previous computational methods using linear techniques with a unique cut-off (35, 36) would introduce a larger error rate. For each subpopulation, scRT profiles were inferred and visualised (Fig. 4E-F). Despite genome-wide CNAs, the pseudo-bulk RT profiles of the 2 MCF-7 subpopulations had a Spearman correlation of 93.6% (Fig. 4G), and also highly correlated with the bulk RT profile (Spearman correlations of 92.8% and 94.4%). As expected, due to the smaller copy-number signatures, the HCT-116 profiles were also highly correlated at 97.5% (Fig. 4H).

**Figure 4.**
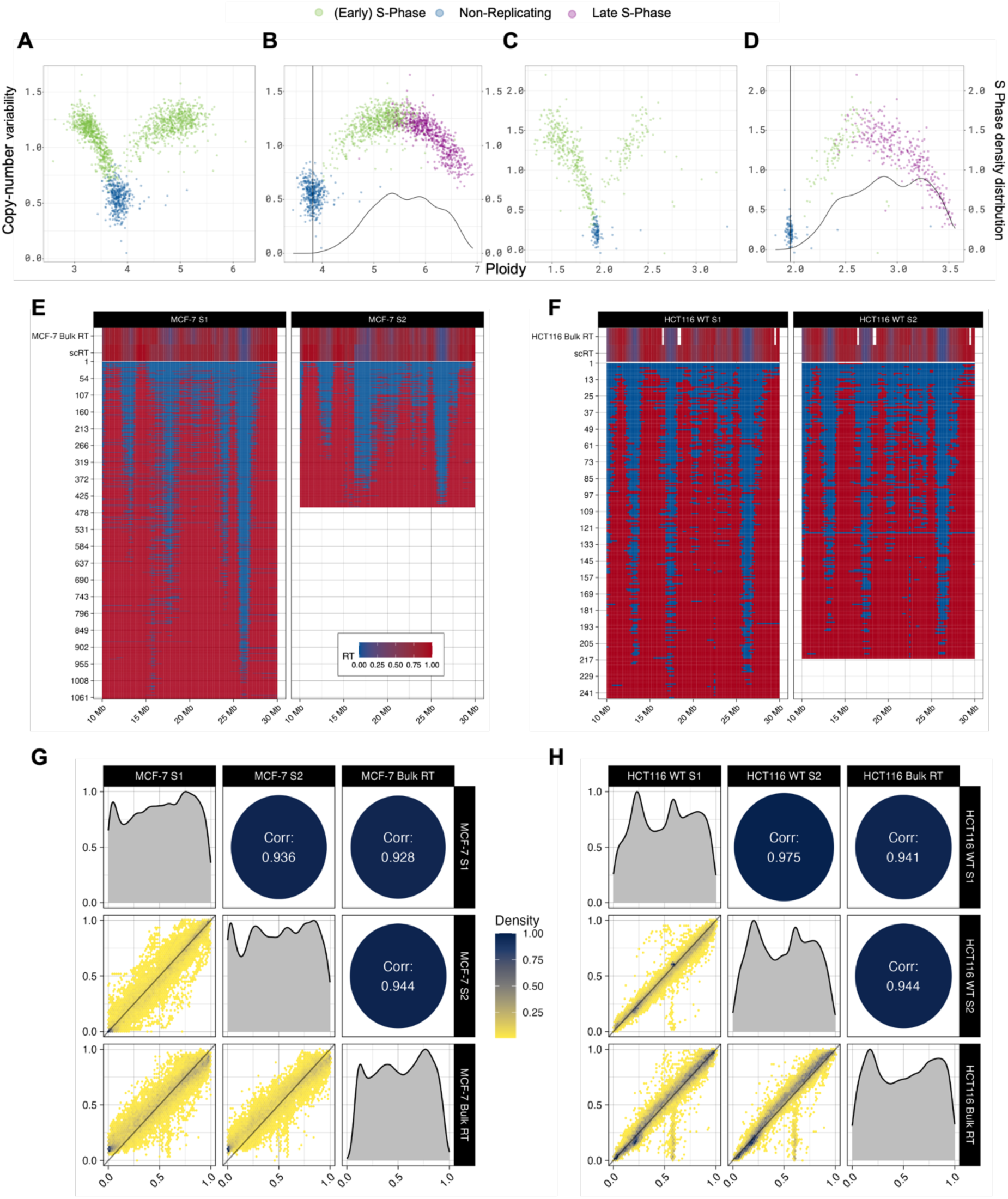
Single-cell Replication Timing (scRT) of heterogenous cancer cell lines uncovered. **(A-D)** Early (green) and late (purple) S-phase cells are corrected (B,D) from raw scCNV data (A,C) displaying non-replicating (blue) and replicating (green) cells. **(E-F)** scRT landscapes of chromosome 16 from MCF-7 (E) and HCT-116 (F) cells display minor differences between subpopulations with pseudo-bulk (scRT) and bulk RT are displayed in the upper window. **(G-H)** Correlations between pseudo-bulk subpopulation scRT and bulk RT for MCF-7 (G) and HCT-116 (H) cells.

### Replication timing changes in patient tumours

Since replication timing in heterogenous tumours has not been studied so far, we used the methods we developed in the same manner as done with the cell-lines to discover cell phases and subpopulations from published data obtained from a triple-negative breast-cancer (TNBC) tumour sample (SA1135). As in the original study (20), we discovered 1 diploid and 3 aneuploid subpopulations from the non-replicating cells (n=58,54,55,26; Fig. 5A-B). Out of the 347 cells that passed quality control, 152 were replicating, showing a larger proportion of replicating cells than those obtained from the cell line models, which is concordant with persistent proliferation of cancer cells. We then calculated the scRT profiles for each subpopulation. We considered that subpopulations 1 (n=13) and 3 (n=29) did not have a representative S phase landscape and disregarded them in the analysis (Fig. 5D). Remarkably, we discovered that subpopulations 2 (n=36) and 4 (n=74) showed distinct replication timing programs, indicating a deregulated replication programme *in vivo*. The two replication profiles from the same tumour correlated at 73.3% demonstrating that RT can be highly modified in subpopulations of the same cancer (Fig. 5C).

**Figure 5.**
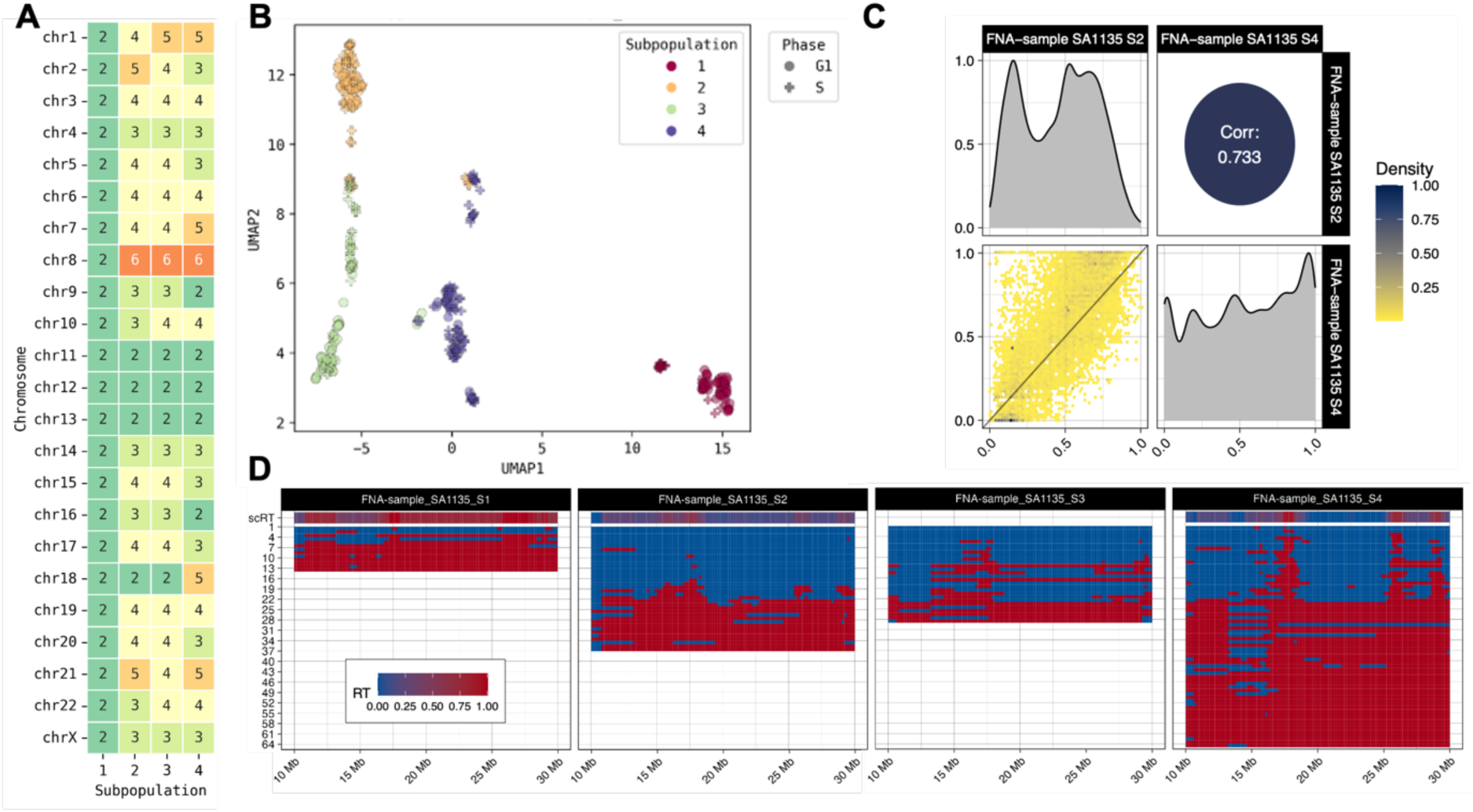
scRT extraction of the subpopulations from the human triple-negative breast cancer sample SA1135. **(A)** median DNA copy-numbers per chromosome for the 4 subpopulations displaying major differences. **(B)** Reduced dimension UMAP plane of the copy-numbers of the single cells of the tumour. **(C)** Spearman correlation between the subpopulations 3 and 4. **(D)** scRT landscapes of chromosome 21 from the 4 subpopulations with the pseudo-bulk RT from the scRT profiles in the upper windows. FNA: Fine-Needle Aspiration (cell collection technique).

### scRT atlas reveals cell-type relationships

We extended the use of our methods to more datasets. In total we analysed the copy-numbers of 119,991 quality-controlled cells originating from 92 different samples spanning across 21 different somatic/cancer cell-lines, 35 patient tumours and 19 patient-derived xenografts (PDX) samples (Supplementary Table 3). In some cases, we discovered large copy-number differences between the same cell lines that were obtained from different publications. HeLa and MCF-7 cell lines displayed a different karyotype depending on their origin, suggesting that extensive culture of cancerous cell lines can induce important copy-number changes (Supplementary Fig. 3). We calculated RT when we had enough cells to reconstruct a representative S phase landscape. This was determined either by software failure or visual inspection of the replication patterns and manual elimination. A total of 41 (sub)populations were used and the Spearman correlation for each pair was calculated (Fig. 6).

**Figure 6.**
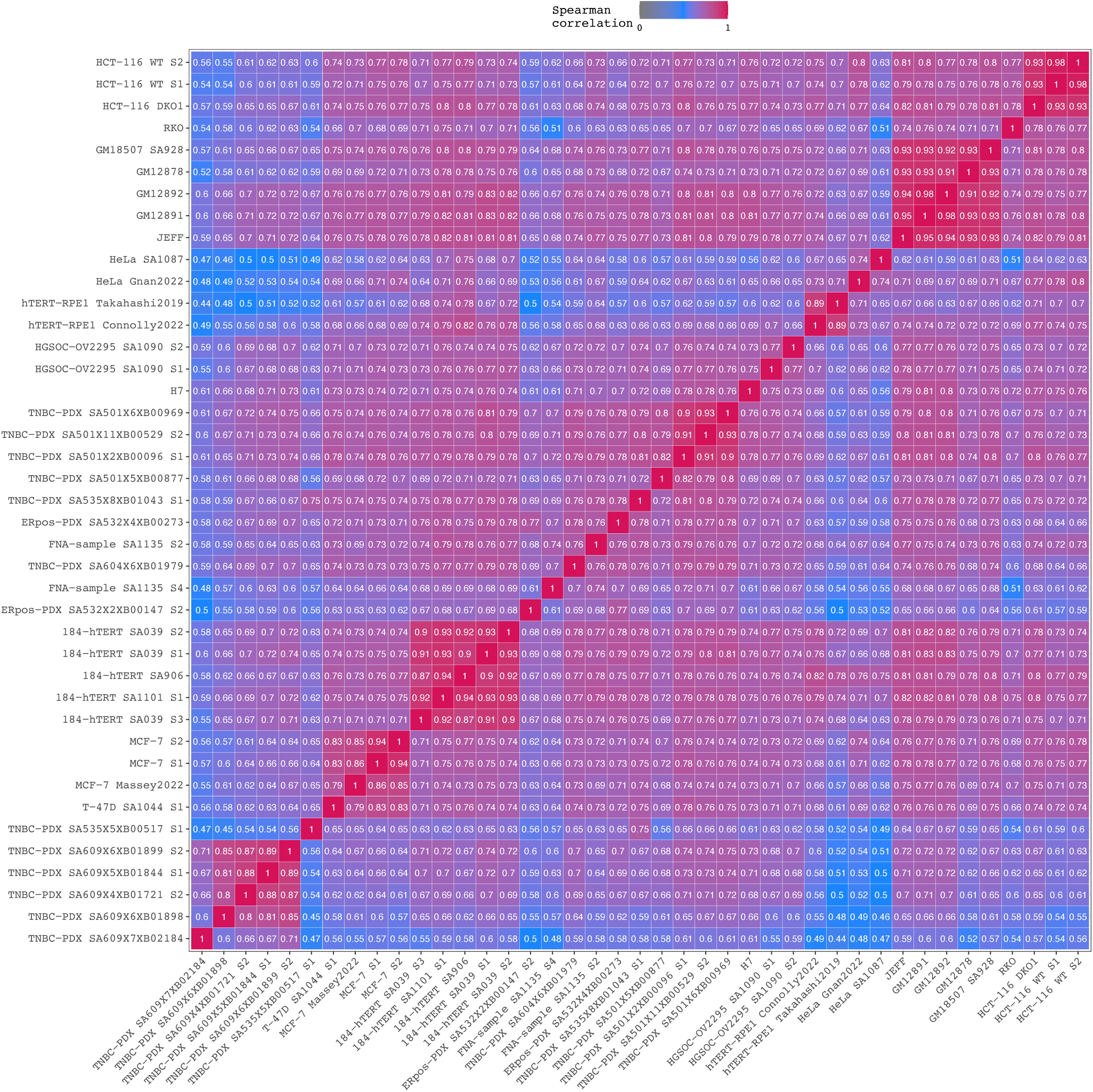
scRT atlas represented by Spearman correlations between the 41 scRT pseudo-bulk profiles extracted from the human (sub)populations in a pan-cancer approach, with samples from cell liens, patient tumors and patient-derived xenografts (PDX) samples. Samples were ordered by hierarchical clustering using the second version of Ward’s minimum variance method (Ward.D2). Subpopulation enumeration is given at the end of the sample name, preceded by S (i.e. S1, S2, etc…). PDX samples are encoded as sample_numberXmouse_passageXsequencing_code. TNBC: Triple-negative breast cancer; ERpos: Estrogen-Receptor positive; PDX: patient-derived xenograft; High-Grade Serous Ovarian Carcinoma; hTERT: human telomerase reverse transcriptase.

In contrast to two subpopulations from the same sample, we noticed that MCF-7 samples from different laboratories only presented an 84.5% correlation, on average. Knowing that this cell line is known to have variable karyotypes, we speculated that the RT differences could be caused by wide-spread copy-number differences. Indeed, we discovered that this cell line only shared between 11 and 13 common median chromosome copies between the two sample origins (Fig. 4, Supplementary Fig. 3). JEFF and GM-lymphoblastoid cell lines on the other hand had extremely high RT correlations all ranging >90%, regardless of the sample origin. Despite both H7 hESCs and GM12892 lymphoblastoid cells presenting a perfectly diploid karyotype (Supplementary Fig 3), their replication tracks presented a 79% correlation, illustrating that in addition to CNV, RT can be used as a cell-type specific biomarker, which can help to determine the potential origin of tumour cells.

### MnM: a fast and accurate tool integrating machine learning for subpopulation discoveries and replication analysis from scWGS data

We integrated our machine learning approaches to provide a single ready-to-use computational tool, MnM – Mix ‘n’ Match, that unifies these techniques under one program. Copy-number imputation, replication state classification and subpopulation detection enabled scRT extraction from heterogenous cell populations and related downstream analyses, for *in vivo* and *in vitro* samples. In addition to the reported accuracies, MnM is a fast tool, with a runtime of 7m:22s for 713 HCT-116 WT cells in 100 kb bins running on a macOS 13.5.2 computer system with 6 intel i5 cores.

## DISCUSSION

In this work, we created a fast and efficient tool to establish single-cell replication states and reveal genomic subpopulations from a single heterogenous sample (such as a tumour sample). MnM uses single-cell copy-numbers from a mixture of heterogenous cells issued from a single sample and discovers heterogeneity to match the replicating and non-replicating cells to their corresponding subpopulations. We individually validated each of the steps that MnM is made of and demonstrated astonishing accuracy for missing-value imputation as well as cell replication state classification. We also confirmed that with subpopulation clustering, MnM can detect the number of different cell-types and as well as the underlying subpopulations from a single sample.

We showed that FACS sorting is prone to a high error rate between 17.7% and 27.7% for cell phase predictions, in accordance with previous estimations (36). Although in some cases there is a legitimate interest to FACS sort single-cells before sequencing, we believe it is essential that the cell-sorted metadata is verified computationally, if possible, to avoid any erroneous conclusions. While our replication state classifier is only trained with the hg38 reference genome, this method can easily be extended to other genomes and be used routinely. Furthermore, in some cases, the FACS metadata may not exists (e.g. unsorted samples). This could be the case of precious samples, such as hESCs and tumours which contain a limited number of cells, one would not want to lose from a restricted yield after FACS sorting. Therefore, *in silico* predictions would be valuable for these cases.

We discovered that JEFF (B lymphocyte) cells had lost a copy of chromosome X, a phenomenon correlated with mitotic errors occurring from ageing (60). We also discovered important chromosomal aberrations in cell-lines and tumour samples (Supplementary Fig. 3) which further underlines the importance of DNA copy-number screening. Aneuploidy is an omnipresent trait in the genomes of tumours (10, 61). Though the presence of genomic instability in cancer has been recognized for a long time (62–64), the exact role it plays in tumour development remains ambiguous. In fact, aneuploidy could act as both a tumour promoter and suppressor (65, 66). As discussed by others (10), copy-number gains might amplify the expression of genes promoting tumours, called oncogenes, cushioned within the altered regions or could stem from the disruption of checkpoint control – a common occurrence in advanced malignancies (67, 68). Nonetheless, aneuploidy might surprisingly exert tumour-suppressive effects in specific cases. For example, genomic instability could decrease tumour fitness (66, 69, 70) while individuals with Down syndrome, arising from the triplication of chromosome 21, exhibit a significantly diminished susceptibility to most solid tumours (71).

The robustness of the DNA replication machinery is a cornerstone of cellular integrity. Without it, it can lead to genomic instability, a hallmark of cancer. In this study, we observed a noteworthy contrast between cell-line models and patient-derived samples in terms of DNA replication timing disruptions. Remarkably, cell-line models exhibited relatively modest distortions in DNA replication dynamics. These models, cultured under controlled conditions, often reflect simplified representations of cellular systems. However, a compelling finding emerged in our analysis of patient samples, where we identified substantial and impactful disruptions in DNA replication patterns. These observations resonate with the idea of the relevance of the tumour microenvironment and its intricate interplay with replication stress and thus genomic stability. The disparities between cell-line and patient sample dynamics highlight the necessity of integrating complex, patient-specific factors into our understanding of DNA replication mechanisms in the context of cancer progression.

Furthermore, whole-genome single-cell studies are still failing to overcome the low coverage reads across the genome. As new methods are emerging with promising advancements, notably a recent report of long-read single-cell sequencing (72), future investigations will be able to dig further into the precise relationship between the mutational landscape, aneuploidy and the replication programme. Eventually, with the imminent generation of higher resolution data, studies will be able to address the replication differences of different homologues with scRT. Thus, we underline the necessity for detailed analyses examining the replication synchronicity of alleles, an even more complex task for aneuploid polyallelic cancers.

In conclusion, we have created a first machine learning based tool to democratise single-cell subpopulation detection from DNA copy-numbers, while also providing a large amount of data building the largest scRT atlas available to date for the community, which could be an important resource for further research. The outcomes of this tool can help contribute to our understanding of cancer emergence and progression. Finally, our results underline the necessity to consider tumour samples in order to fully understand the mechanisms governing DNA replication in cancer. Although cell lines constitute an easier research model to study, they lack some critical environmental factors that interact with cancer.

## DATA AVAILABILITY

The source code of MnM are available at the GitHub page of the team (https://github.com/CL-CHEN-Lab/MnM) and was registered at French Agency for the Protection of Programs (APP) under registration number N° IDDN.FR.001.340005.000.S.P.2023.000.31230. The 10x barcode whitelist can be found at https://github.com/TheKorenLab/Single-cell-replication-timing/blob/main/align/10x_barcode_whitelist.txt. The R script to discover qualitative barcodes from single-cells through the expectation-maximisation algorithm, the python script to split subpopulation and replicate copy-number files, related code and scCNV matrices from the data can be found at the MnM GitHub depository as well. The human reference genome hg38 can be found at https://support.illumina.com/sequencing/sequencing_software/igenome.html. The genomic blacklist can be found at: https://github.com/Boyle-Lab/Blacklist. Kronos scRT can be found at https://github.com/CL-CHEN-Lab/Kronos_scRT and the modified version of Kronos used here can be found at https://github.com/josephides/Kronos_scRT. Published single-cell datasets can be found on the Gene Expression Omnibus (GEO) under the accession numbers <u>GSE186173</u> (35), <u>GSE158011</u> (30), <u>GSE108556</u> (33), on the Sequence Read Archive (SRA) under <u>PRJNA770772</u> (36) and on the European Genome-Phenome archive (EGA) under <u>EGAS00001003190</u> (20). Processed scCNV data from (24) can be found at https://zenodo.org/record/6998936. Bulk RT profiles were obtained under accession numbers <u>GSE34399</u> for MCF-7 and <u>GSE158011</u> for HCT-116 cells while the liftover chain file is available at https://hgdownload.cse.ucsc.edu/goldenpath/hg19/liftOver.

## SUPPLEMENTARY DATA

Supplementary Data including Supplementary Figures and Supplementary Tables are available together with the manuscript.

## AUTHOR CONTRIBUTIONS

C.L.C. conceived and planned the study. J.M.J. developed the program and performed the bioinformatics analyses. C.L.C. supervised the development and bioinformatics analyses. J.M.J. and C.L.C. wrote the manuscript.

## ACKNOWLEDGEMENTS

Part of this manuscript was prepared using a limited access dataset obtained from BC Cancer and does not necessarily reflect the opinions or views BC Cancer. The authors would like to acknowledge Tatiana Popova and Guillem Rigaill for stimulating discussions, Jean-Baptiste Guillaumin and Emeline Ravalli for the APP application as well as Yoann Schumacher for establishing the program’s license.

