## Supplemental Figures for "MnM: a machine learning approach to detect replication states and genomic subpopulations for single-cell DNA replication timing disentanglement"

Supplementary Figures

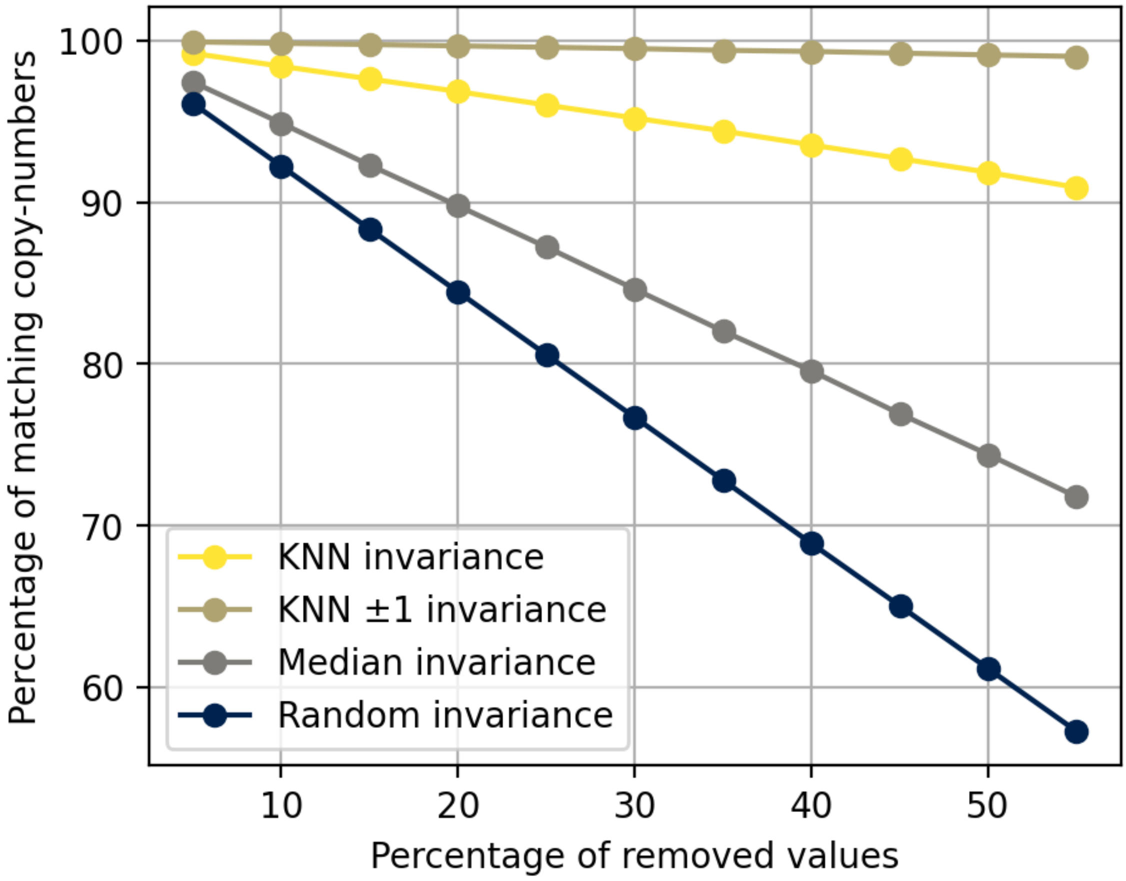


**Supplementary Figure S1.** KNN imputation is an efficient method compensating for single-cell copy-number scarcity. Missing values were simulated by using MCF-7 copy-numbers (data from Gnan et al. 2022) in 100 kb windows which underwent random value removal ranging from 5 to 55% of the total number of values in the single-cell copy-number matrix (regions/cells). KNN, median and random imputations were performed while KNN imputed values that varied by ±1 copy-number were calculated. These four metrics were compared to the original values for copy-number matrix-wide invariance. Four simulations with different seeds were performed to obtain the mean and standard deviation (SD) values for each percentage of removed values with the KNN accuracy. However, the SD values (ranging from 0.024 to 0.0819, average 0.0456) were too small to be visible on the plot.

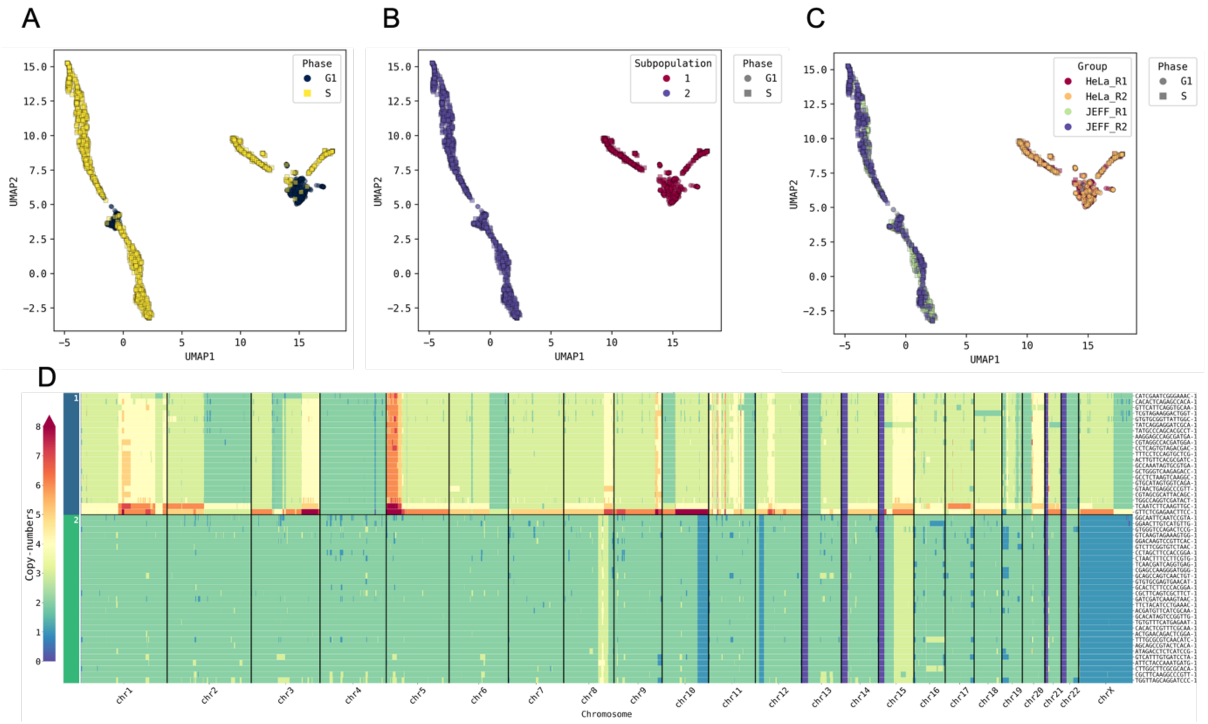


**Supplementary Figure S2.** scCNV distinctions with unsupervised learning. A-C. UMAP pane of JEFF and HeLa samples coloured by replication state **(A)**, subpopulation **(B)** and replicate state of final detected groups **(C)**. **(D)** 50 randomly selected cells and their genome-wide single-cell copy-numbers.

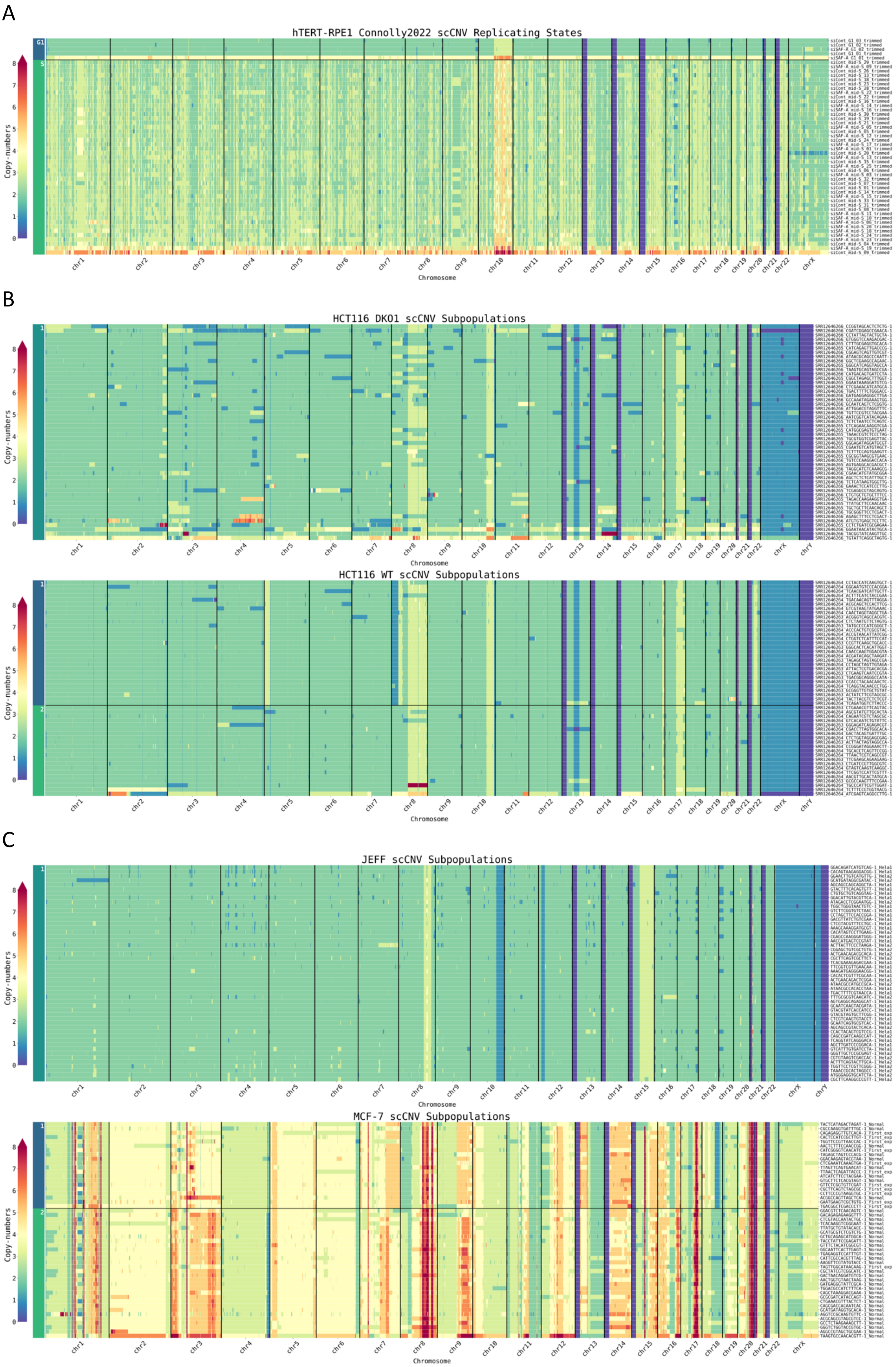
A
hTERT-RPE1 Connolly2022 scCNV Replicating States
B
HCT116 DKO1 scCNV Subpopulations
HCT116 WT scCNV Subpopulations
C
JEFF scCNV Subpopulations
MCF-7 scCNV Subpopulations
Copy-numbers
Chromosome

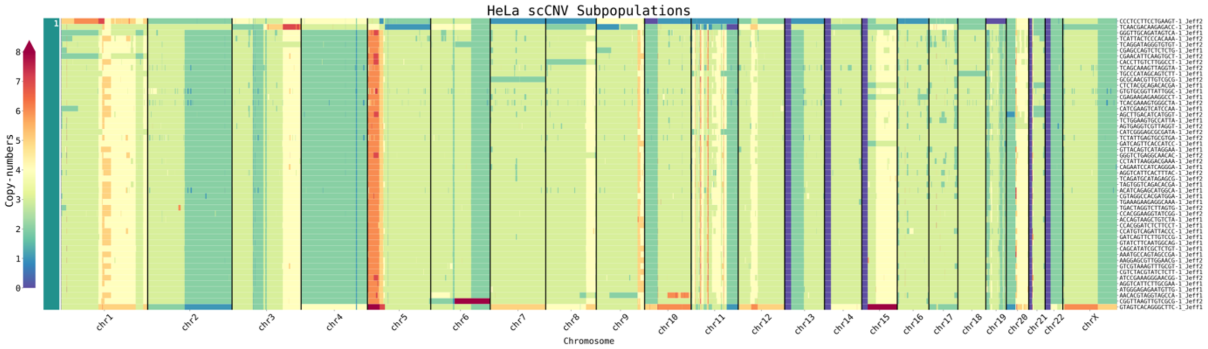
HeLa scCNV Subpopulations
Copy-numbers
Chromosome
chr1 chr2 chr3 chr4 chr5 chr6 chr7 chr8 chr9 chr10 chr11 chr12 chr13 chr14 chr15 chr16 chr17 chr18 chr19 chr20 chr21 chr22 chrX

D

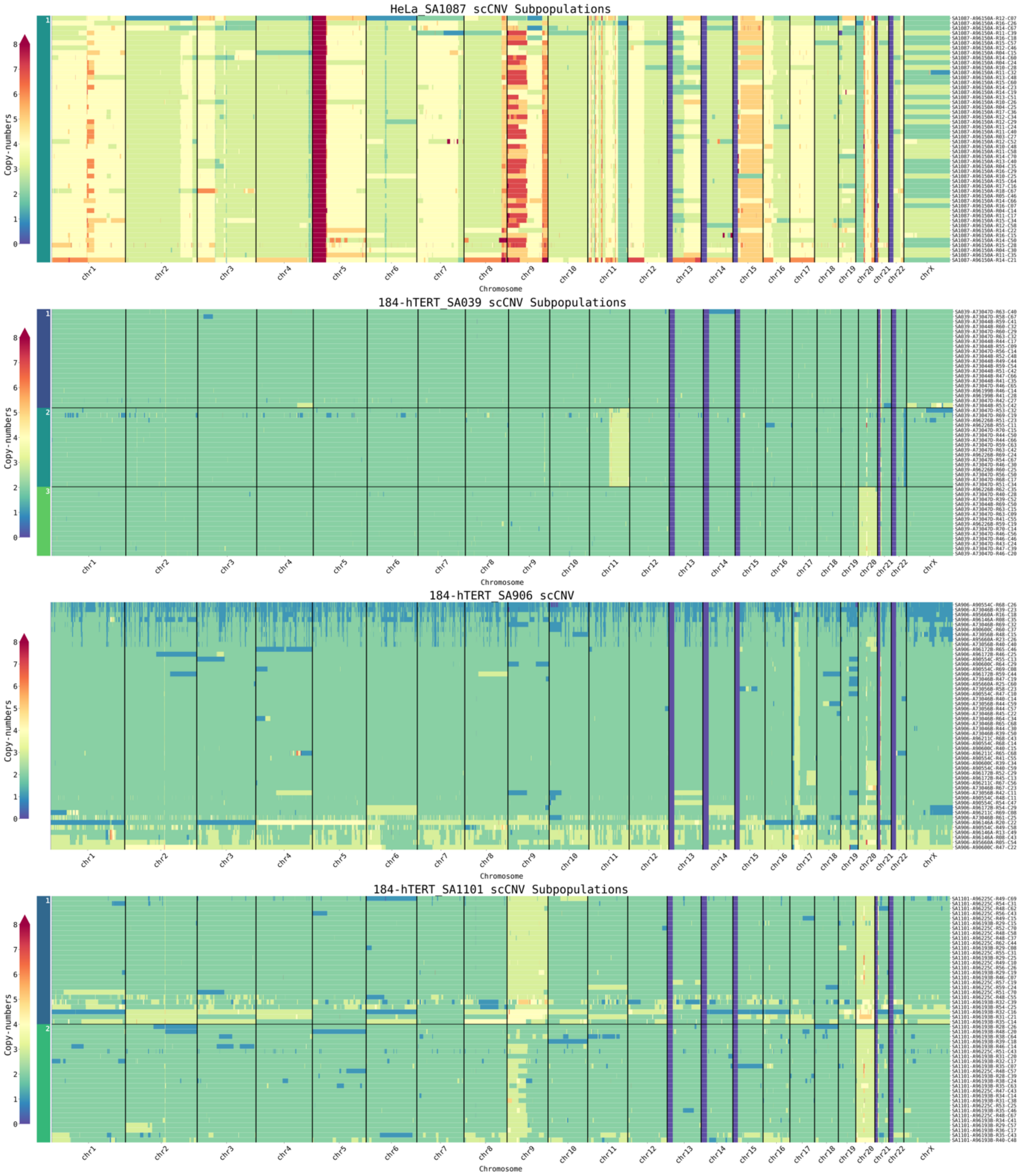
HeLa_SA1087 scCNV Subpopulations
184-hTERT_SA039 scCNV Subpopulations
184-hTERT_SA906 scCNV
184-hTERT_SA1101 scCNV Subpopulations
Copy-numbers
Chromosome
chr1 chr2 chr3 chr4 chr5 chr6 chr7 chr8 chr9 chr10 chr11 chr12 chr13 chr14 chr15 chr16 chr17 chr18 chr19 chr20 chr21 chr22 chrX

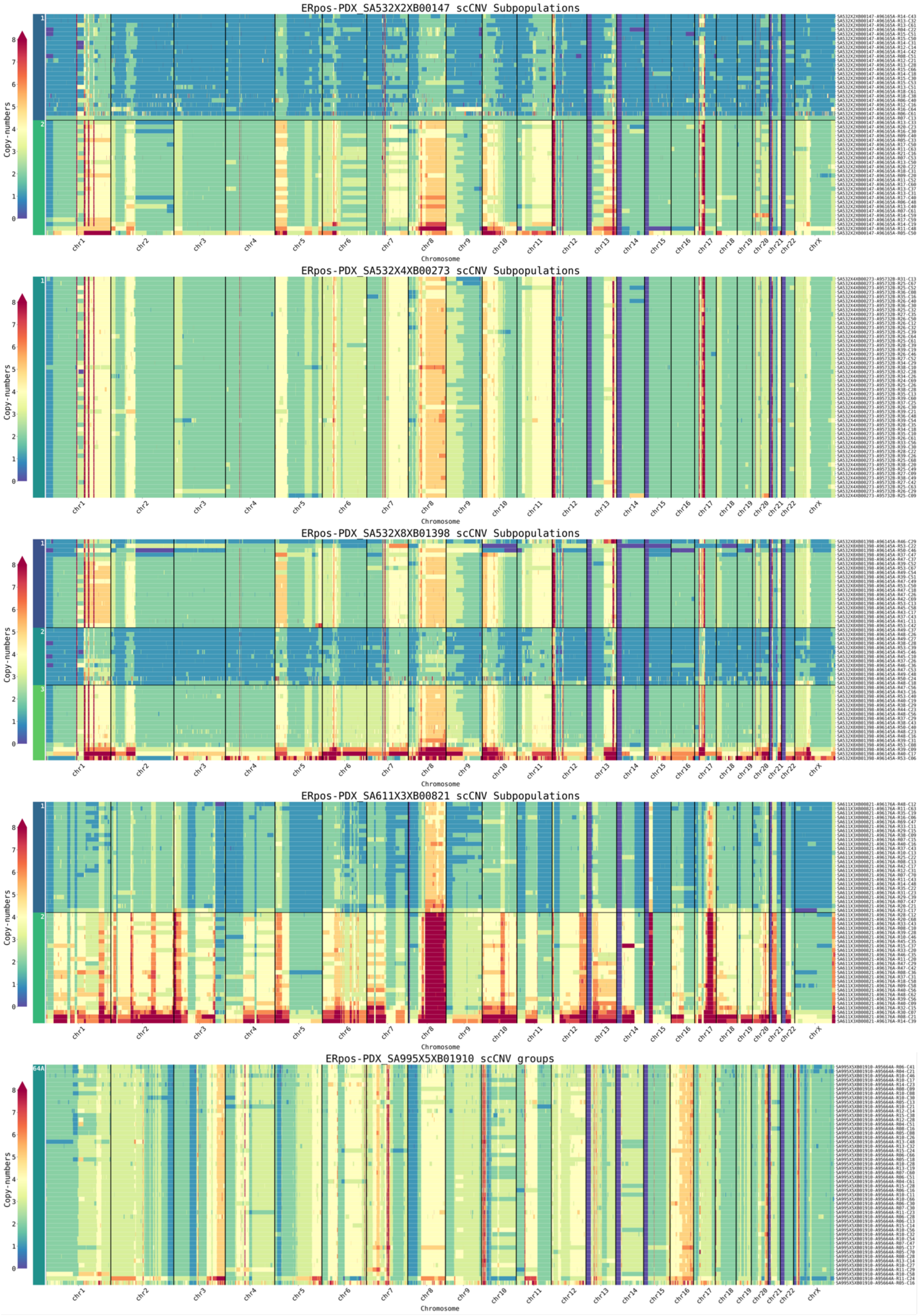
ERpos-PDX_SA532X2XB00147 scCNV Subpopulations
ERpos-PDX_SA532X4XB00273 scCNV Subpopulations
ERpos-PDX_SA532X8XB01398 scCNV Subpopulations
ERpos-PDX_SA611X3XB00821 scCNV Subpopulations
ERpos-PDX_SA995X5XB01910 scCNV groups
Copy-numbers
0
1
2
3
4
5
6
7
8
chr1 chr2 chr3 chr4 chr5 chr6 chr7 chr8 chr9 chr10 chr11 chr12 chr13 chr14 chr15 chr16 chr17 chr18 chr19 chr20 chr21 chr22 chrX
Chromosome

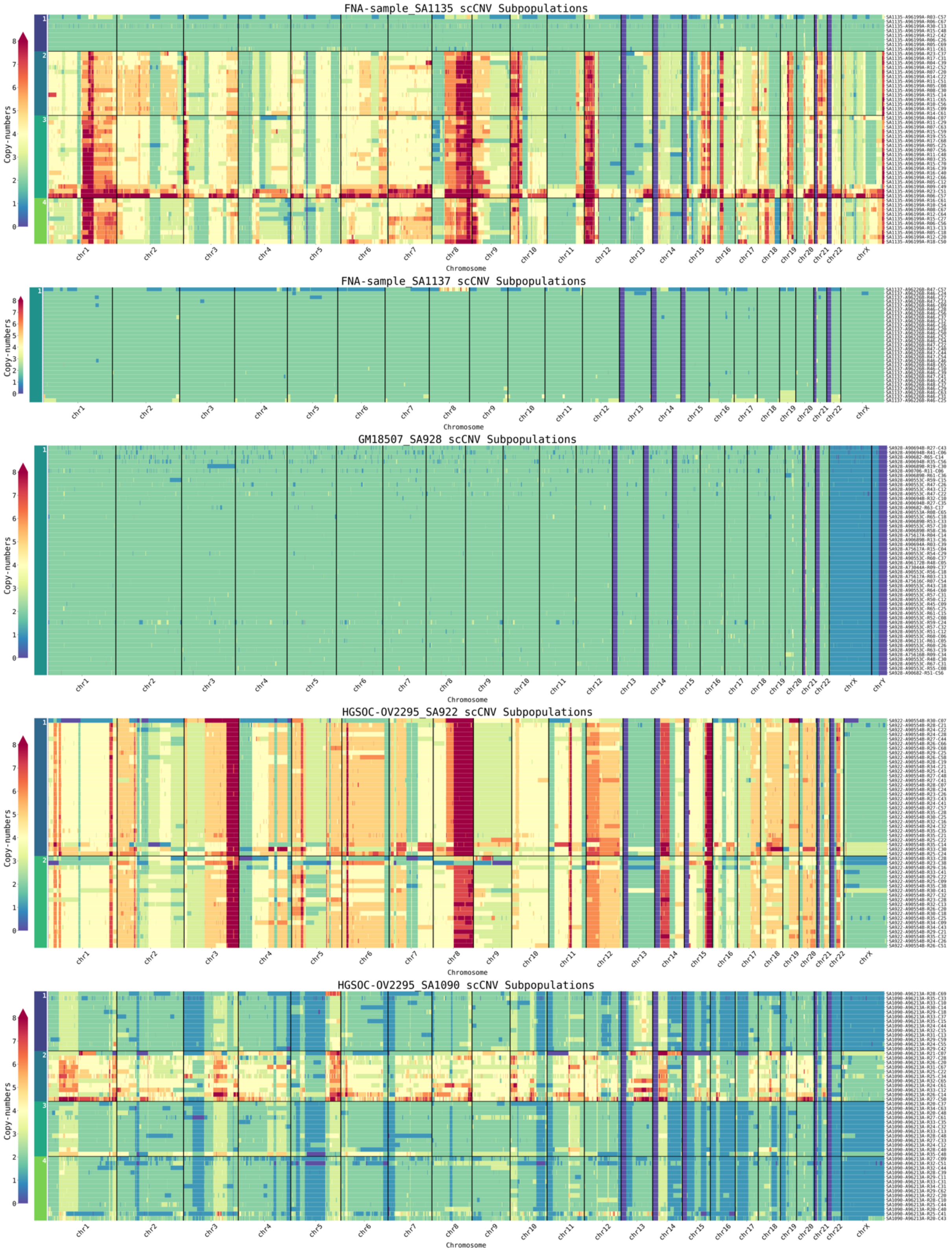

FNA-sample_SA1135 scCNV Subpopulations
FNA-sample_SA1137 scCNV Subpopulations
GM18507_SA928 scCNV Subpopulations
HGSOC-OV2295_SA922 scCNV Subpopulations
HGSOC-OV2295_SA1090 scCNV Subpopulations
Copy-numbers
Chromosome
chr1 chr2 chr3 chr4 chr5 chr6 chr7 chr8 chr9 chr10 chr11 chr12 chr13 chr14 chr15 chr16 chr17 chr18 chr19 chr20 chr21 chr22 chrX
chrY

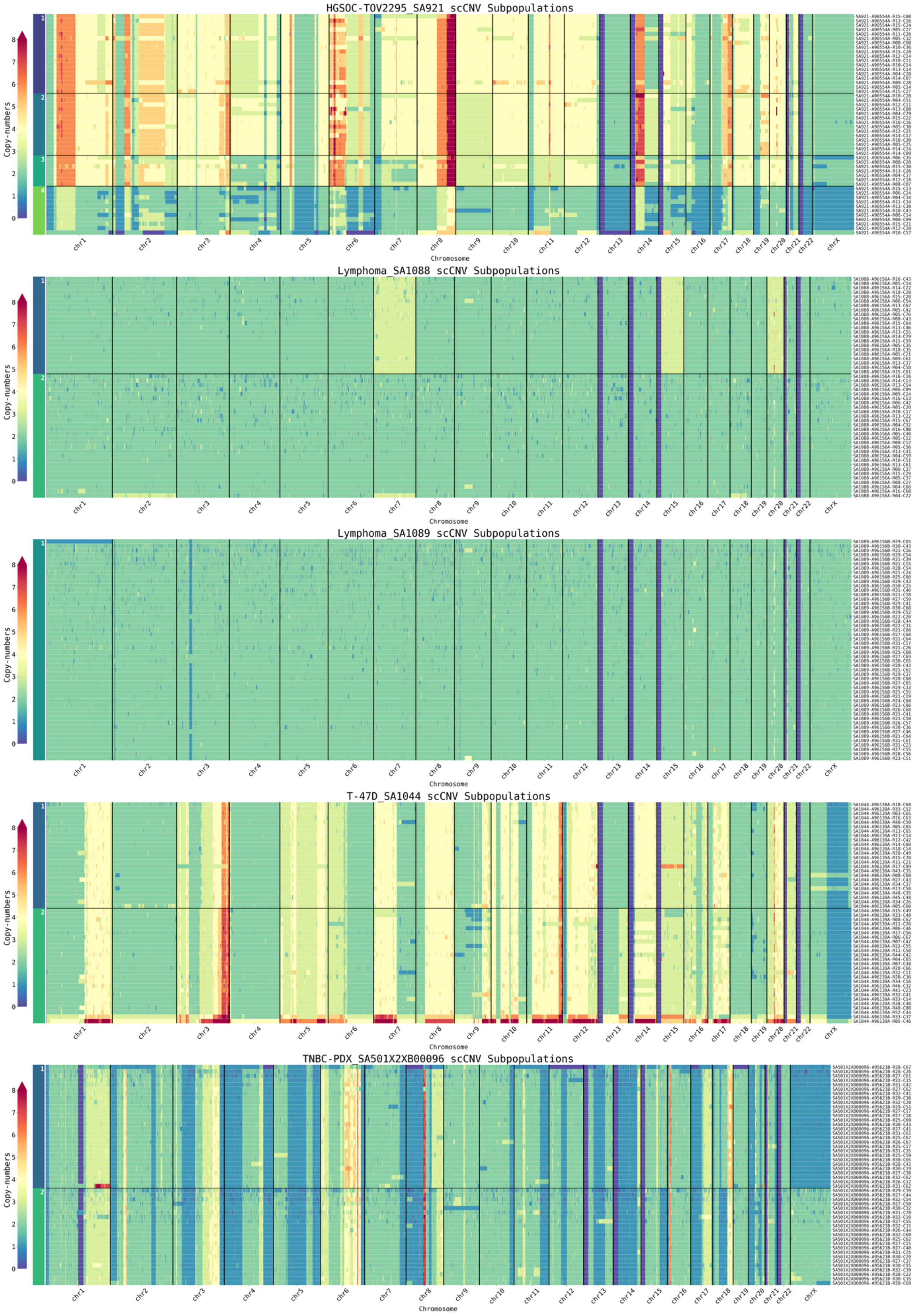
HGSOC-TOV2295_SA921 scCNV Subpopulations
Lymphoma_SA1088 scCNV Subpopulations
Lymphoma_SA1089 scCNV Subpopulations
T-47D_SA1044 scCNV Subpopulations
TNBC-PDX_SA501X2XB00096 scCNV Subpopulations
Copy-numbers
0
1
2
3
4
5
6
7
8
chr1
chr2
chr3
chr4
chr5
chr6
chr7
chr8
chr9
chr10
chr11
chr12
chr13
chr14
chr15
chr16
chr17
chr18
chr19
chr20
chr21
chr22
chrX
Chromosome

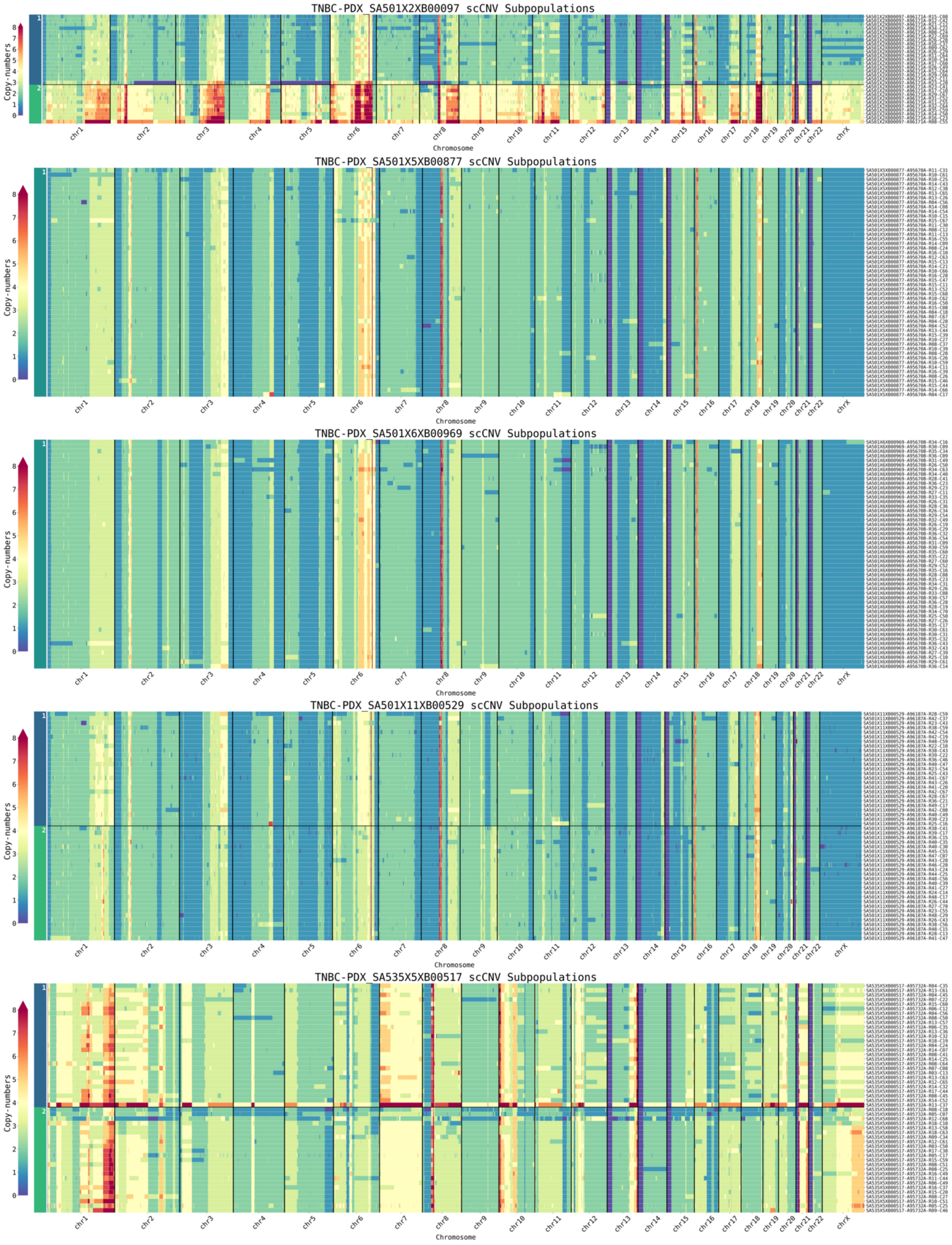
TNBC-PDX_SA501X2XB00097 scCNV Subpopulations
TNBC-PDX_SA501X5XB00877 scCNV Subpopulations
TNBC-PDX_SA501X6XB00969 scCNV Subpopulations
TNBC-PDX_SA501X11XB00529 scCNV Subpopulations
TNBC-PDX_SA535X5XB00517 scCNV Subpopulations
Copy-numbers
Chromosome
chr1
chr2
chr3
chr4
chr5
chr6
chr7
chr8
chr9
chr10
chr11
chr12
chr13
chr14
chr15
chr16
chr17
chr18
chr19
chr20
chr21
chr22
chrX

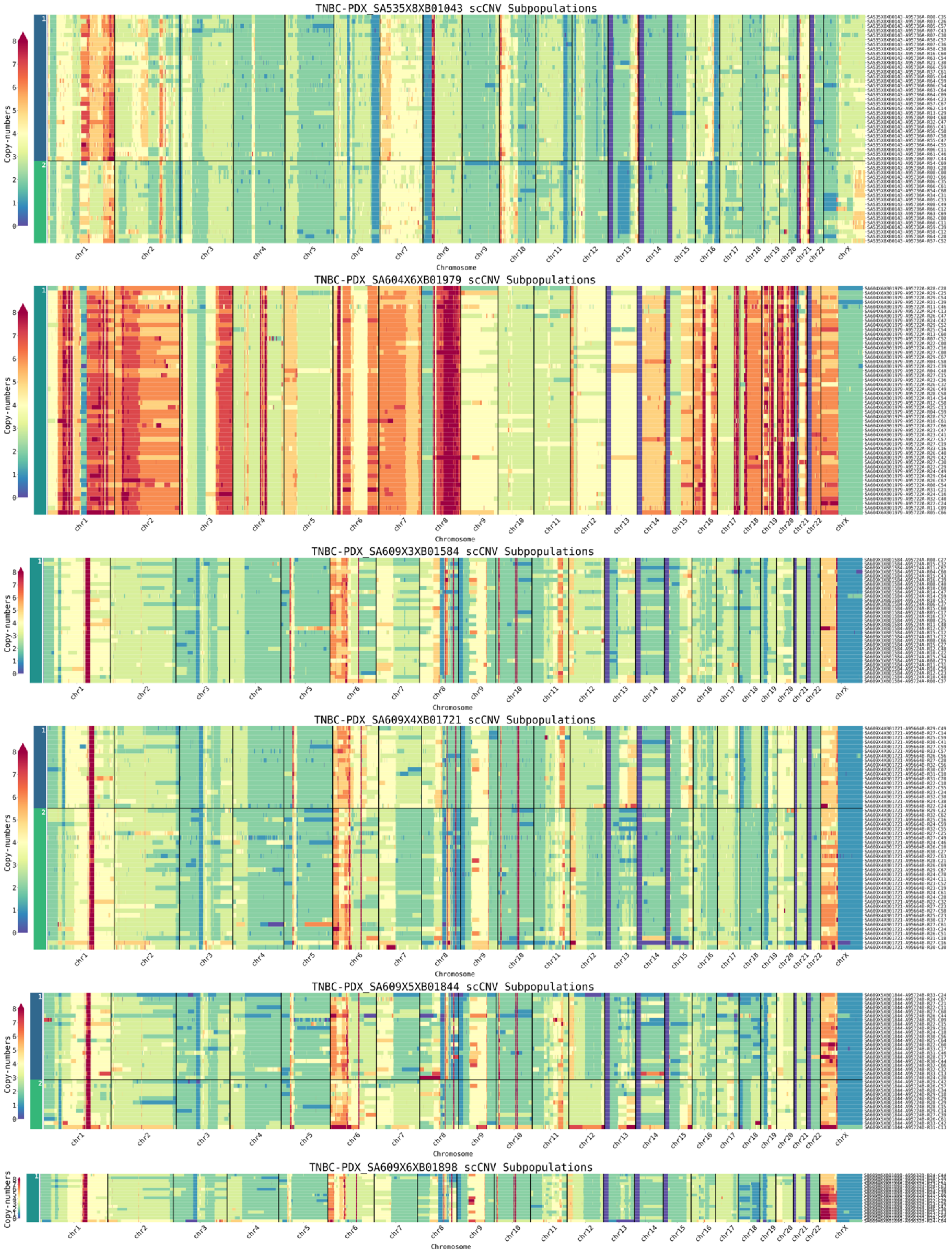
TNBC-PDX_SA535X8XB01043 scCNV Subpopulations
TNBC-PDX_SA604X6XB01979 scCNV Subpopulations
TNBC-PDX_SA609X3XB01584 scCNV Subpopulations
TNBC-PDX_SA609X4XB01721 scCNV Subpopulations
TNBC-PDX_SA609X5XB01844 scCNV Subpopulations
TNBC-PDX_SA609X6XB01898 scCNV Subpopulations
Copy-numbers
0 1 2 3 4 5 6 7 8
chr1 chr2 chr3 chr4 chr5 chr6 chr7 chr8 chr9 chr10 chr11 chr12 chr13 chr14 chr15 chr16 chr17 chr18 chr19 chr20 chr21 chr22 chrX
Chromosome

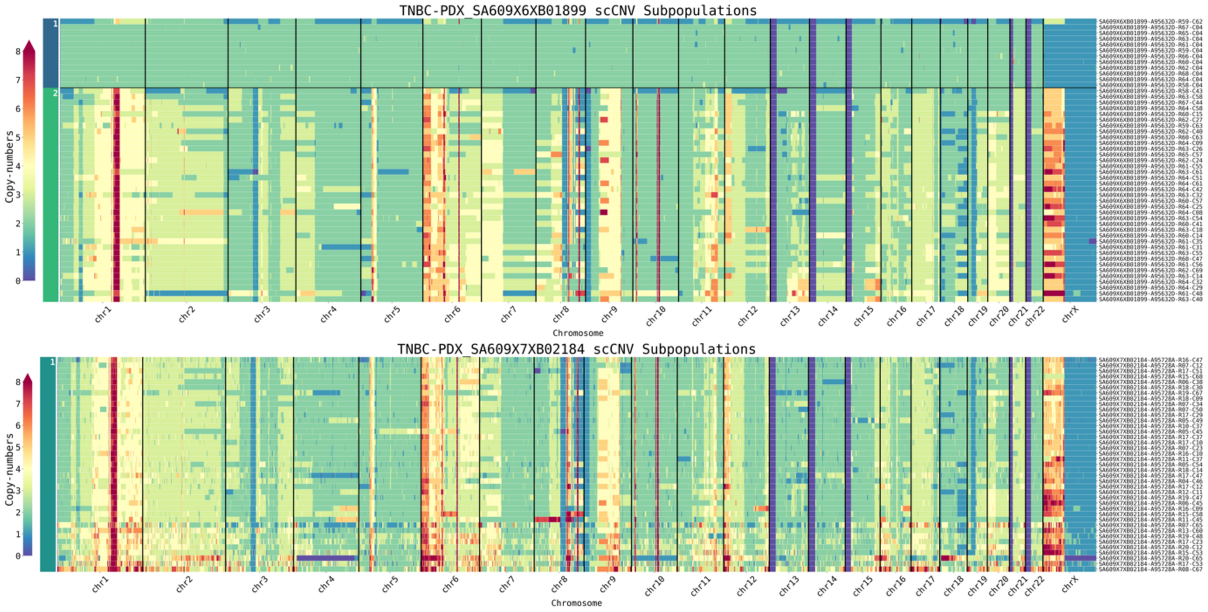
TNBC-PDX_SA609X6XB01899 scCNV Subpopulations
Copy-numbers
Chromosome
TNBC-PDX_SA609X7XB02184 scCNV Subpopulations
Copy-numbers
Chromosome

E

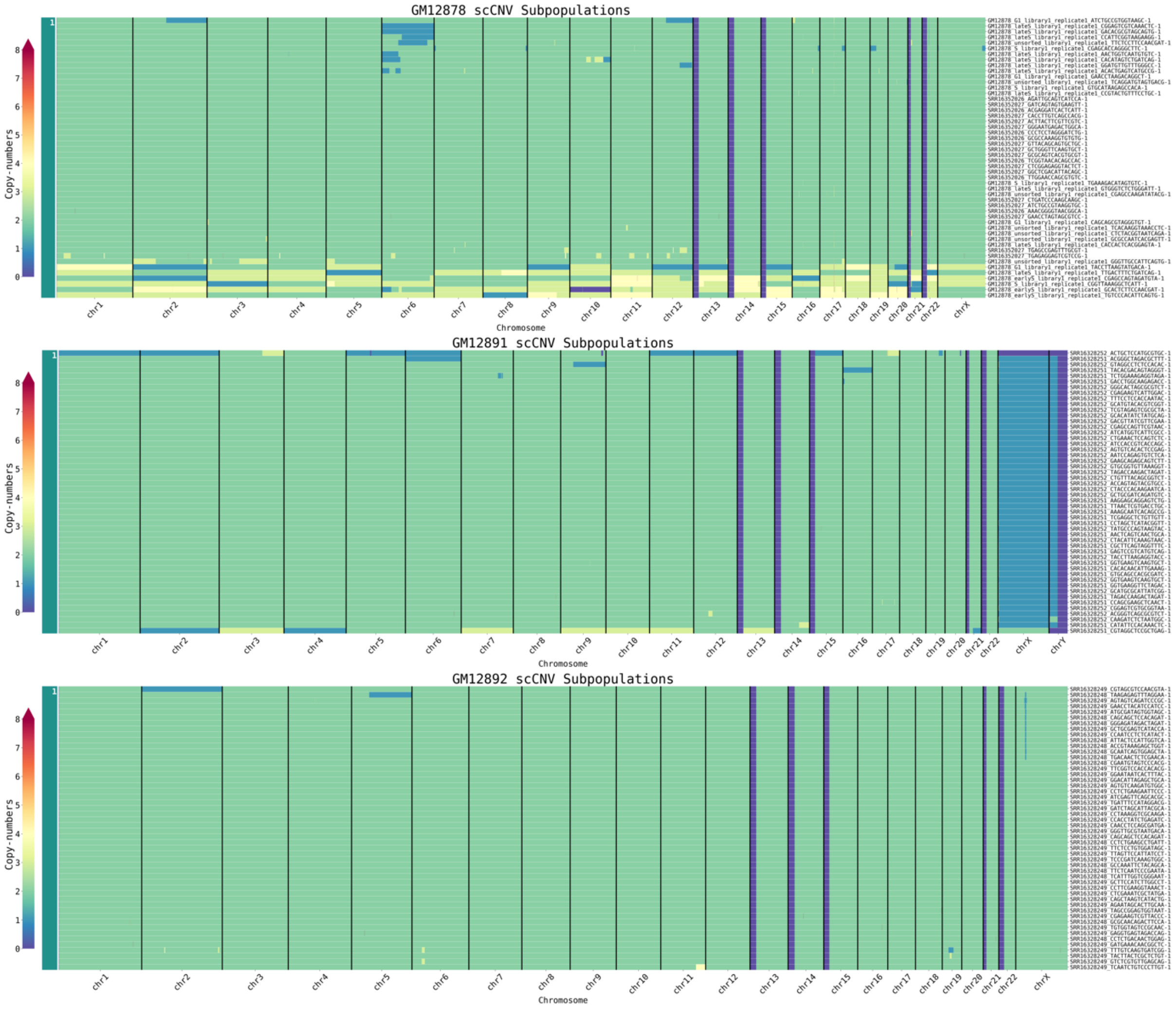
GM12878 scCNV Subpopulations
Copy-numbers
Chromosome
GM12891 scCNV Subpopulations
Copy-numbers
Chromosome
GM12892 scCNV Subpopulations
Copy-numbers
Chromosome

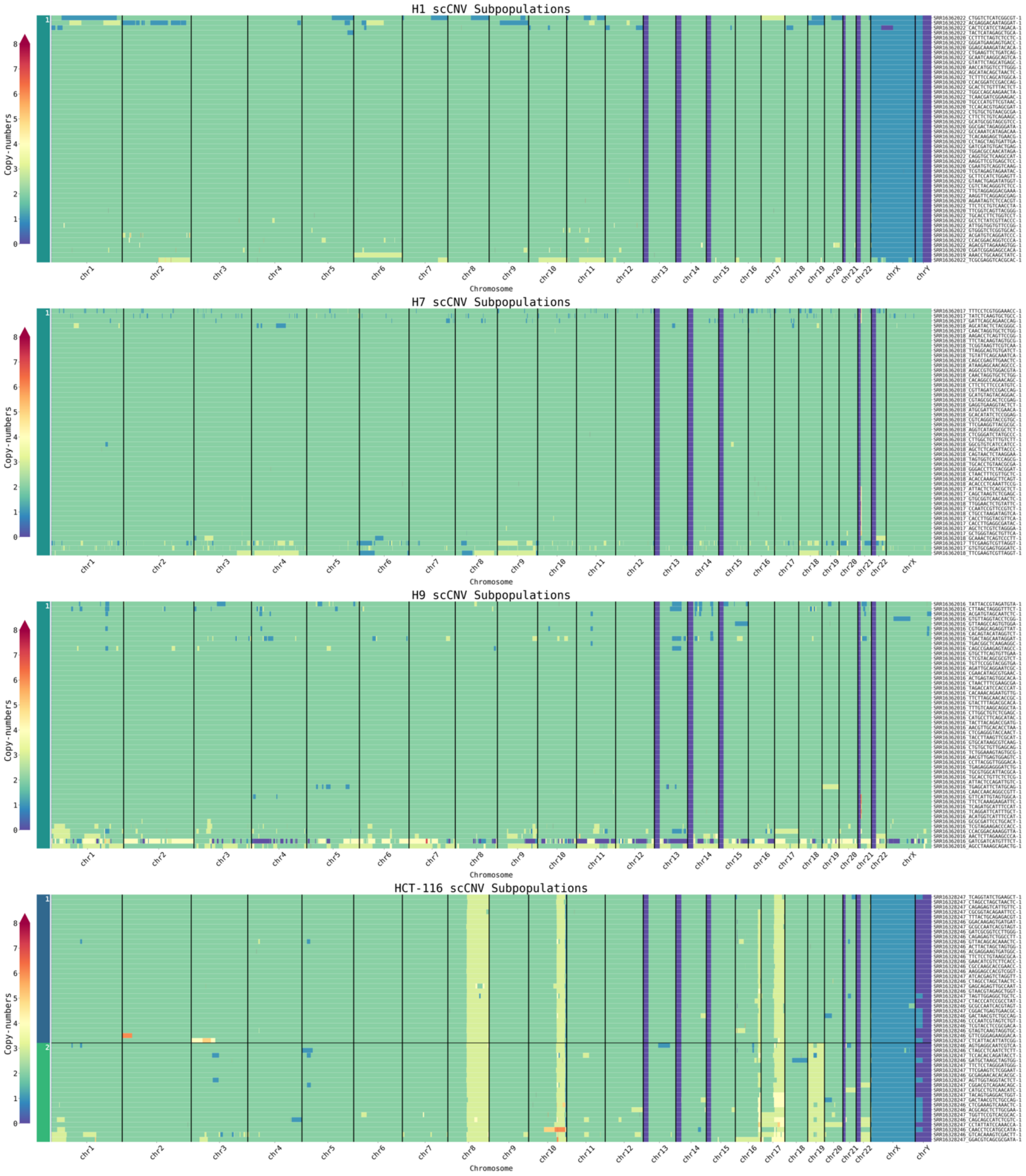
H1 scCNV Subpopulations
H7 scCNV Subpopulations
H9 scCNV Subpopulations
HCT-116 scCNV Subpopulations
Copy-numbers
0
1
2
3
4
5
6
7
8
Chromosome
chr1
chr2
chr3
chr4
chr5
chr6
chr7
chr8
chr9
chr10
chr11
chr12
chr13
chr14
chr15
chr16
chr17
chr18
chr19
chr20
chr21
chr22
chrX
chrY

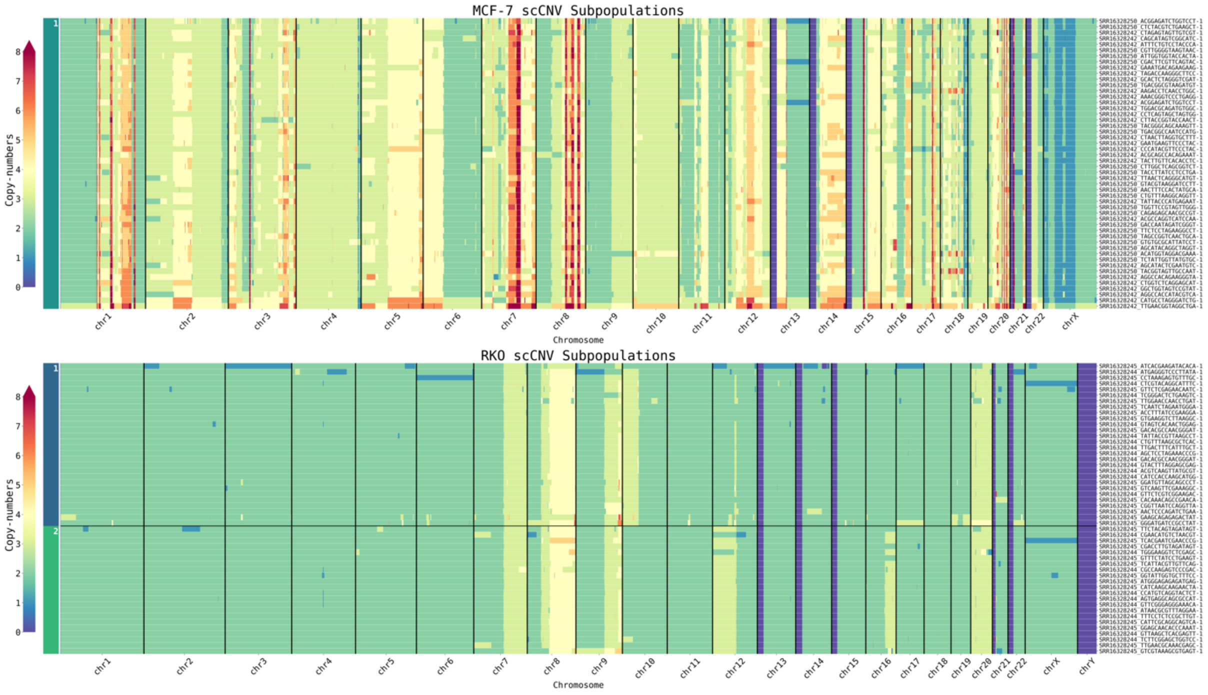
MCF-7 scCNV Subpopulations
Copy-numbers
Chromosome
RKO scCNV Subpopulations
Copy-numbers
Chromosome

F

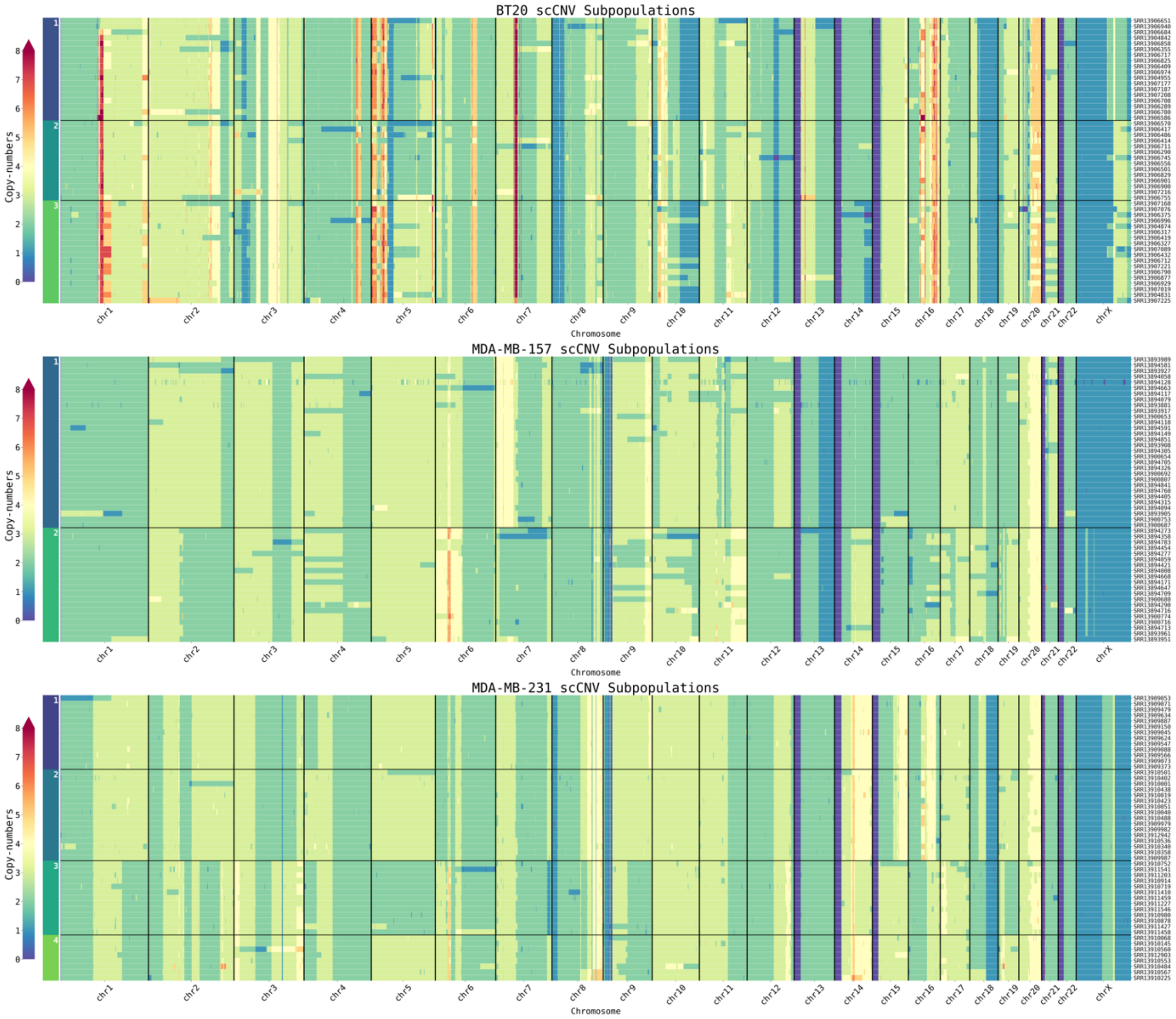
BT20 scCNV Subpopulations
Copy-numbers
Chromosome
MDA-MB-157 scCNV Subpopulations
Copy-numbers
Chromosome
MDA-MB-231 scCNV Subpopulations
Copy-numbers
Chromosome

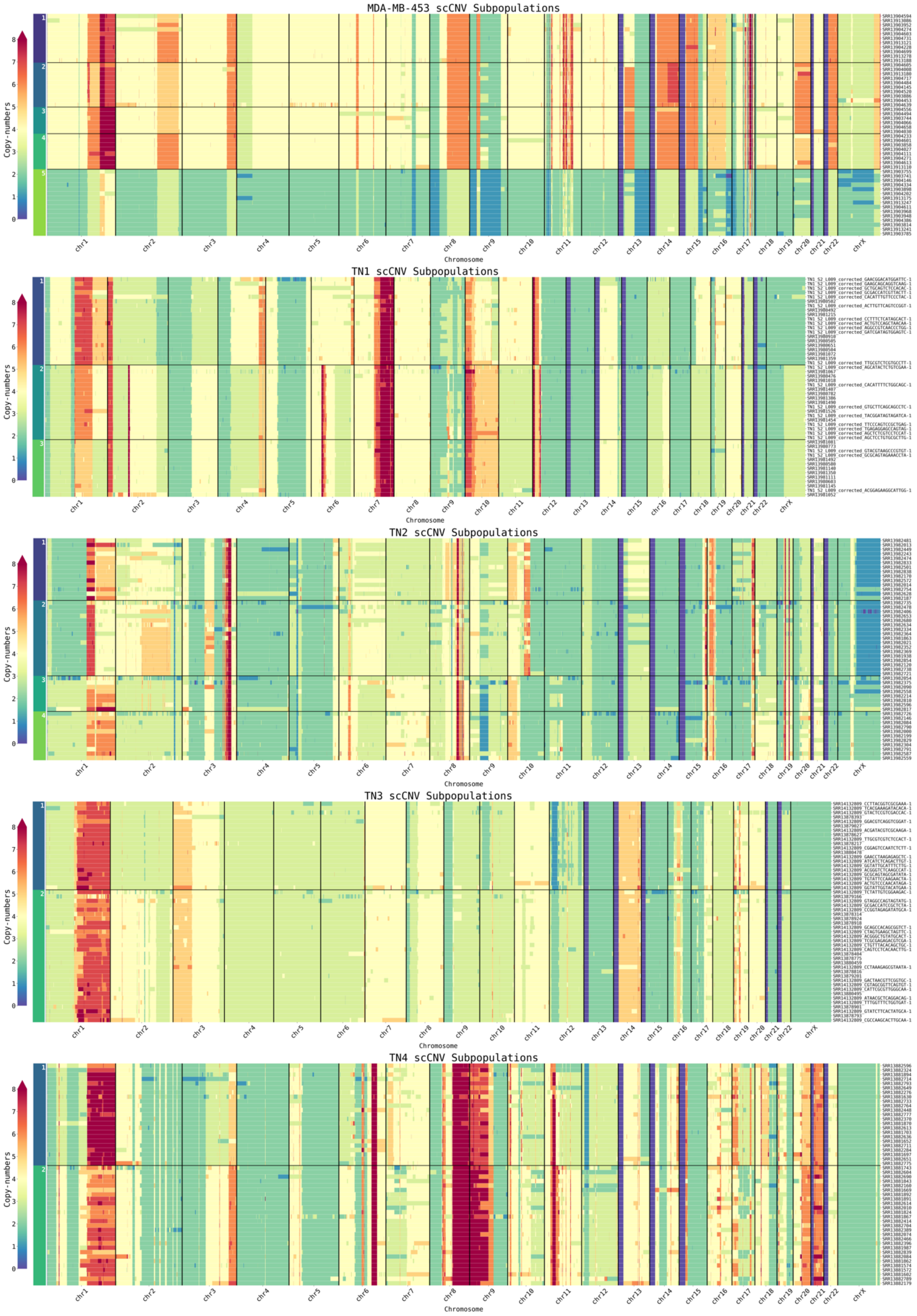

MDA-MB-453 scCNV Subpopulations
TN1 scCNV Subpopulations
TN2 scCNV Subpopulations
TN3 scCNV Subpopulations
TN4 scCNV Subpopulations
Copy-numbers
0
1
2
3
4
5
6
7
8
chr1
chr2
chr3
chr4
chr5
chr6
chr7
chr8
chr9
chr10
chr11
chr12
chr13
chr14
chr15
chr16
chr17
chr18
chr19
chr20
chr21
chr22
chrX
Chromosome

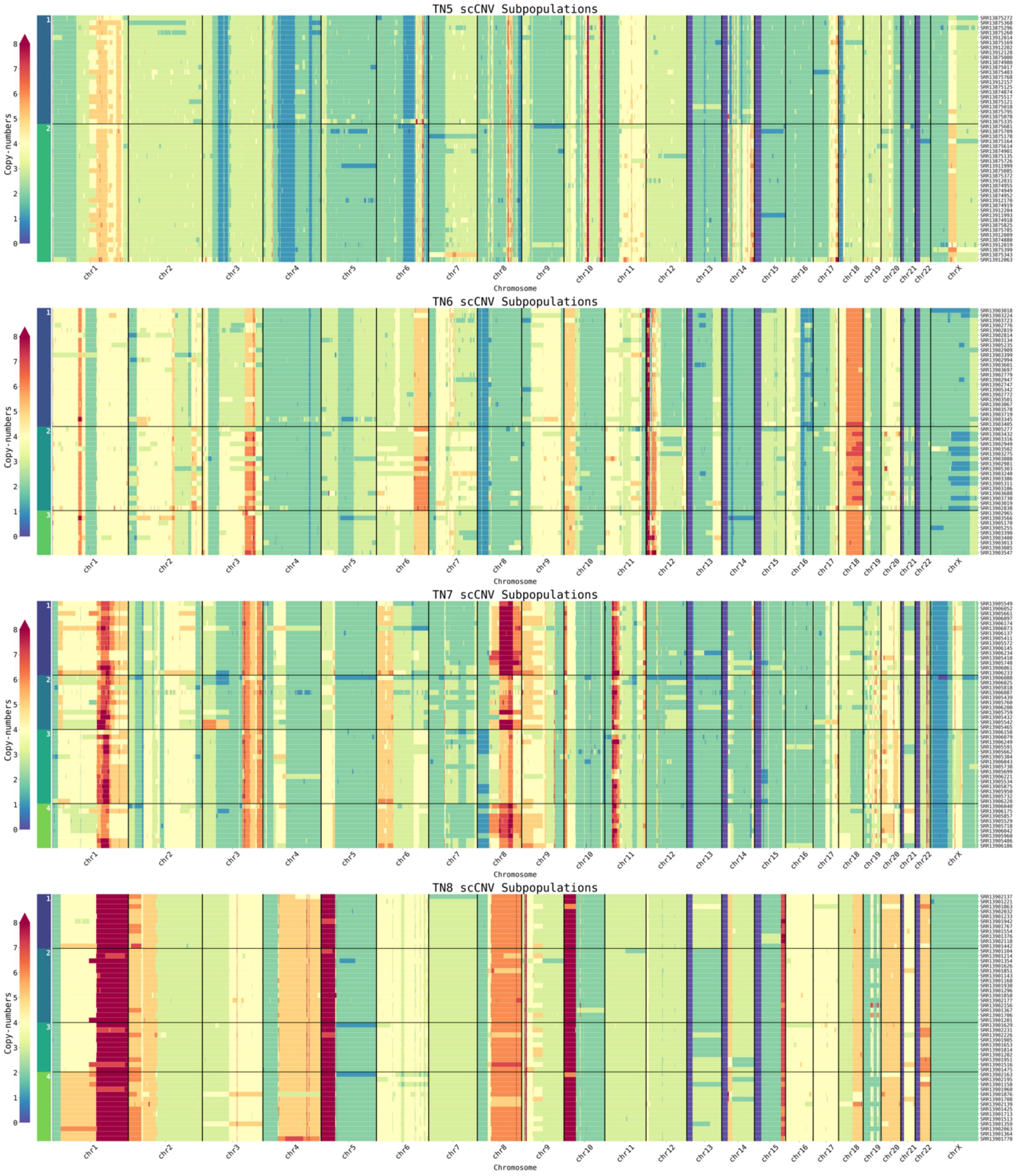
TN5 scCNV Subpopulations
TN6 scCNV Subpopulations
TN7 scCNV Subpopulations
TN8 scCNV Subpopulations
Copy-numbers
Chromosome
chr1
chr2
chr3
chr4
chr5
chr6
chr7
chr8
chr9
chr10
chr11
chr12
chr13
chr14
chr15
chr16
chr17
chr18
chr19
chr20
chr21
chr22
chrX

G

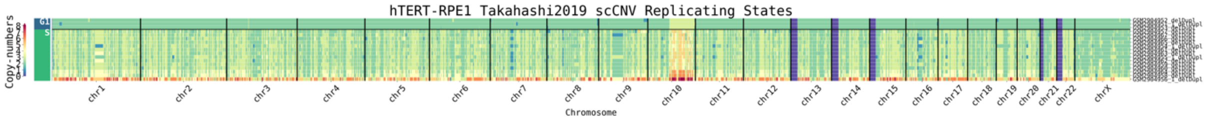
hTERT-RPE1 Takahashi2019 scCNV Replicating States
Copy-numbers
Chromosome
chr1
chr2
chr3
chr4
chr5
chr6
chr7
chr8
chr9
chr10
chr11
chr12
chr13
chr14
chr15
chr16
chr17
chr18
chr19
chr20
chr21
chr22
chrX

H

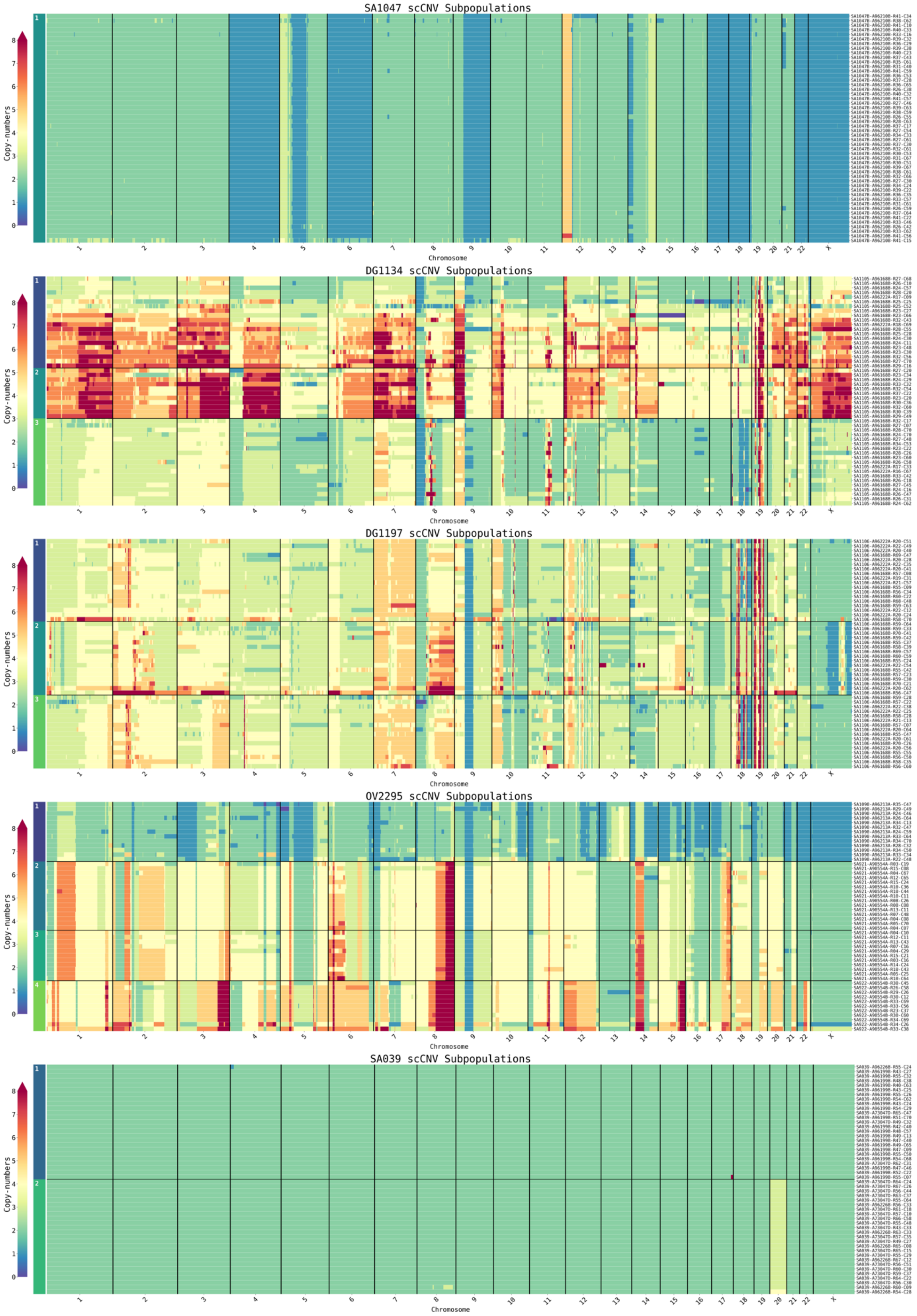
SA1047 scCNV Subpopulations
DG1134 scCNV Subpopulations
DG1197 scCNV Subpopulations
OV2295 scCNV Subpopulations
SA039 scCNV Subpopulations
Copy-numbers
Chromosome

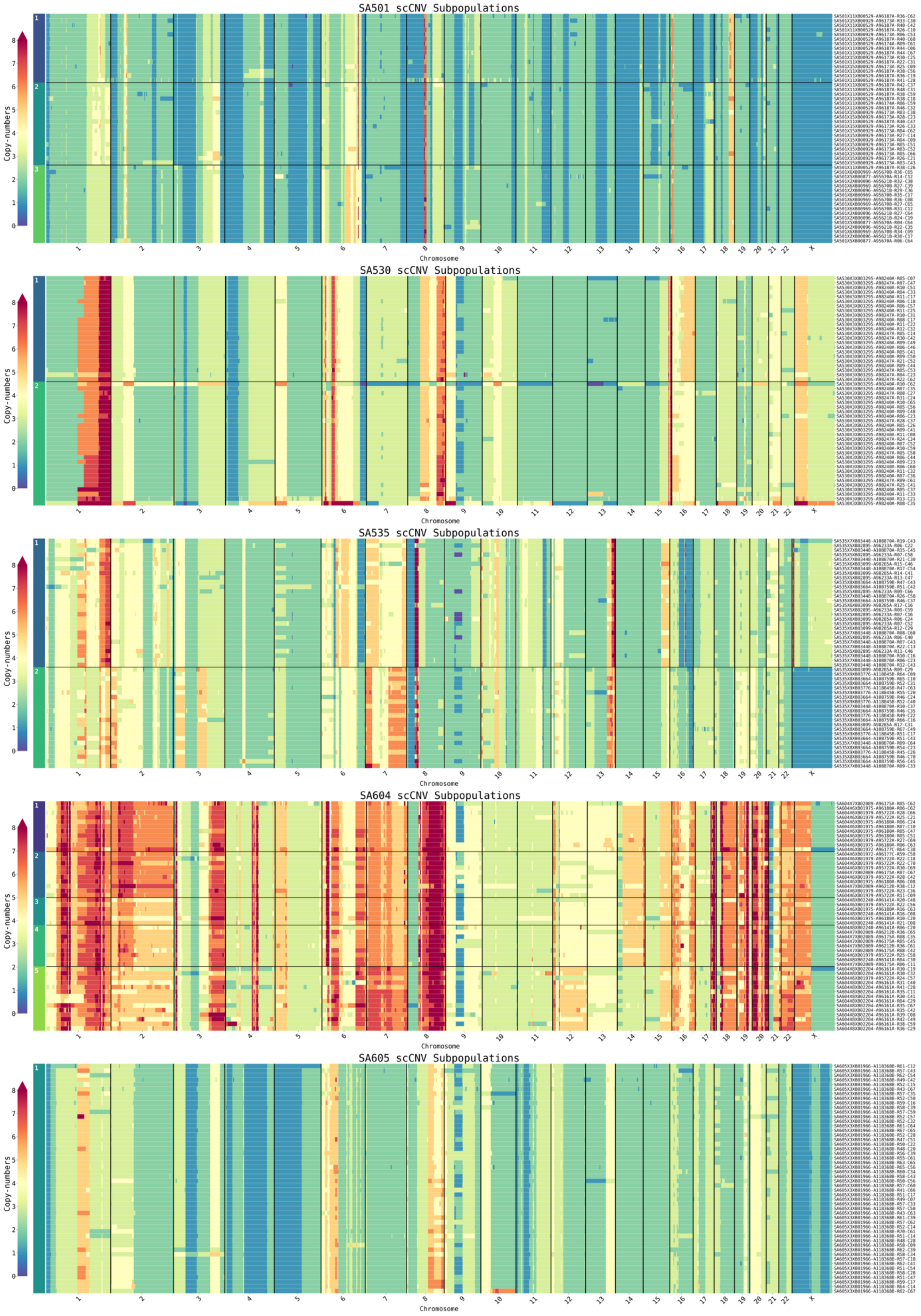
SA501 scCNV Subpopulations
SA530 scCNV Subpopulations
SA535 scCNV Subpopulations
SA604 scCNV Subpopulations
SA605 scCNV Subpopulations
Copy-numbers
Chromosome

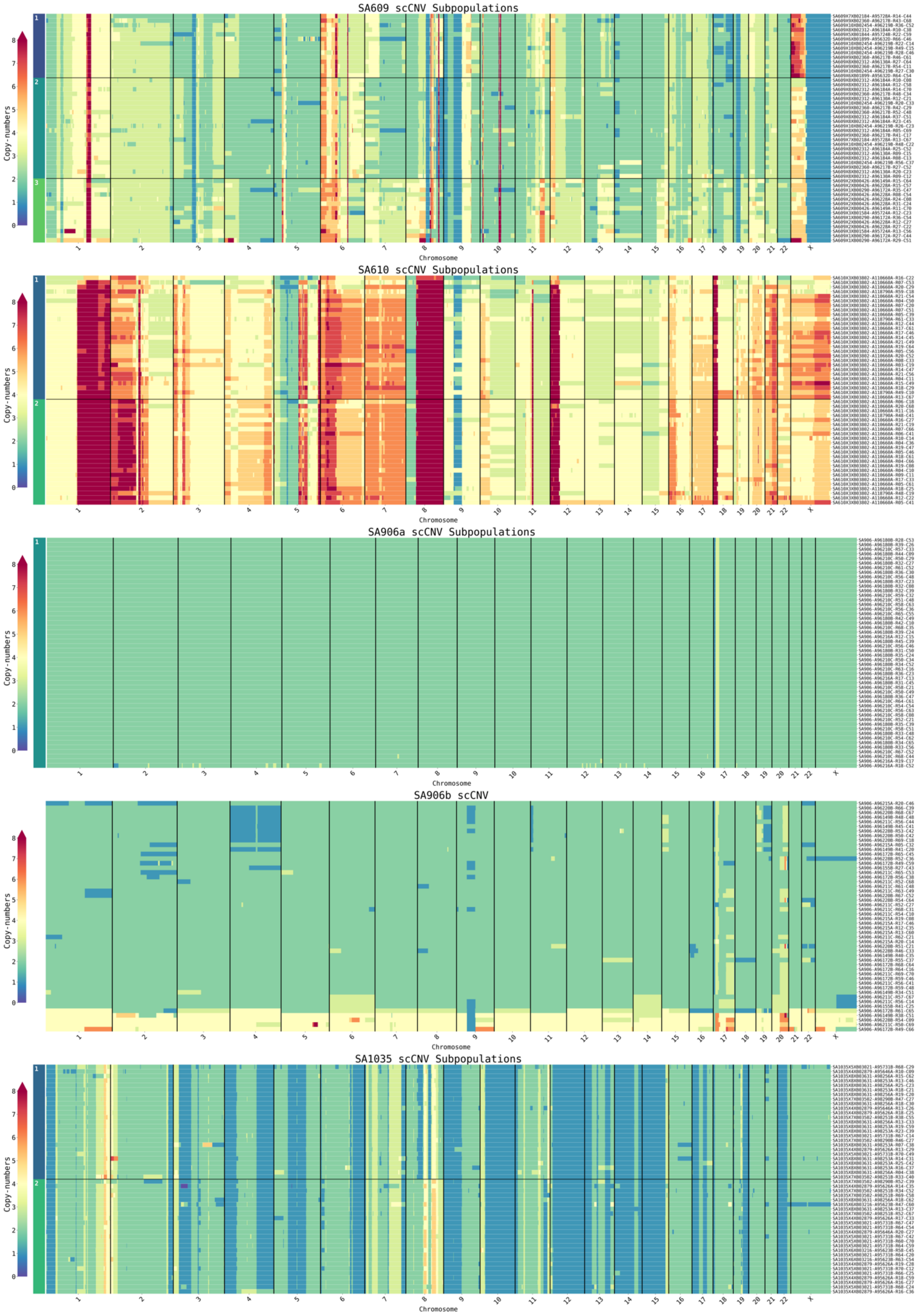

SA609 scCNV Subpopulations
SA610 scCNV Subpopulations
SA906a scCNV Subpopulations
SA906b scCNV
SA1035 scCNV Subpopulations
Copy-numbers
Chromosome

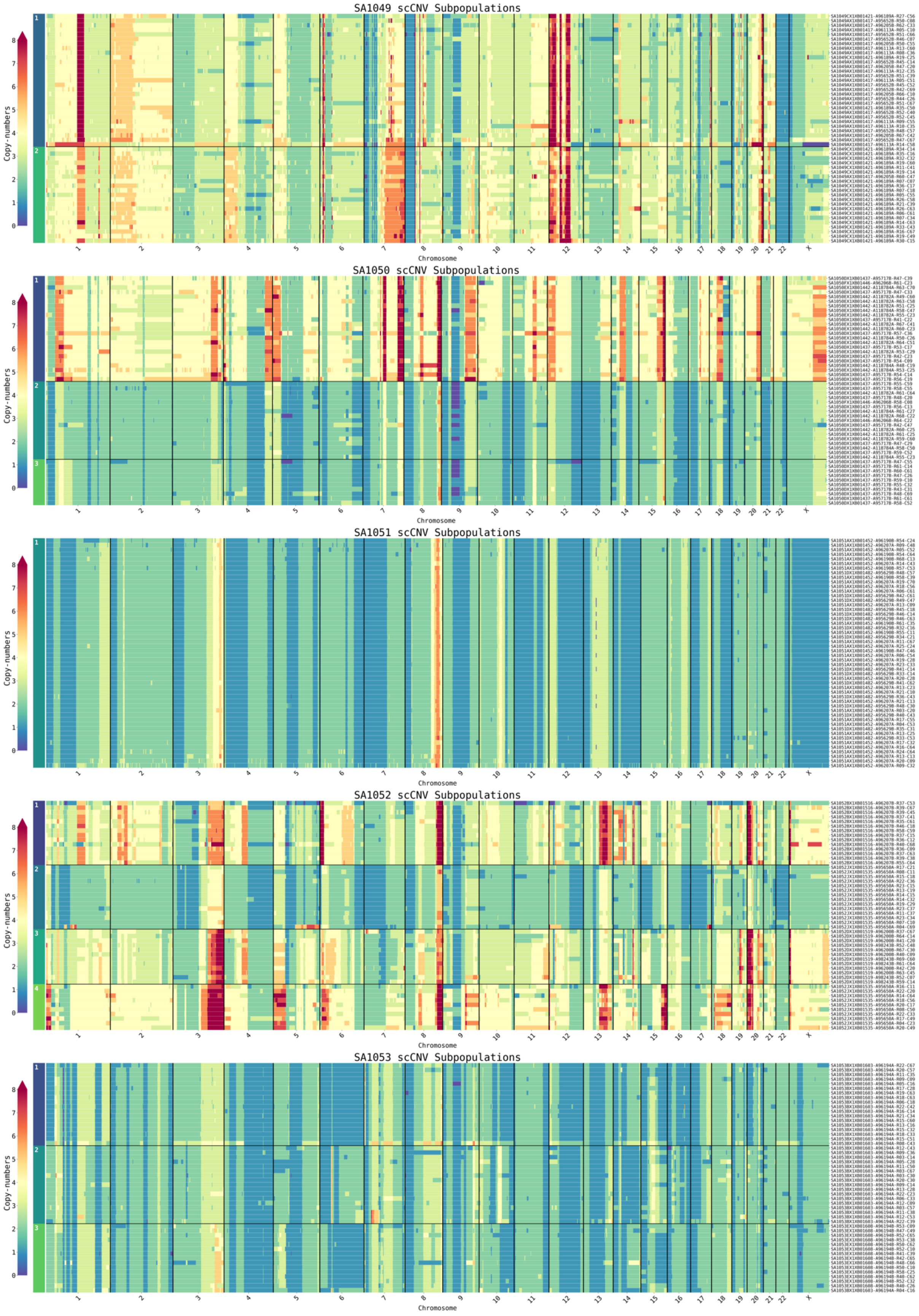
SA1049 scCNV Subpopulations
SA1050 scCNV Subpopulations
SA1051 scCNV Subpopulations
SA1052 scCNV Subpopulations
SA1053 scCNV Subpopulations
Copy-numbers
Chromosome
1 2 3 4 5 6 7 8 9 10 11 12 13 14 15 16 17 18 19 20 21 22 X

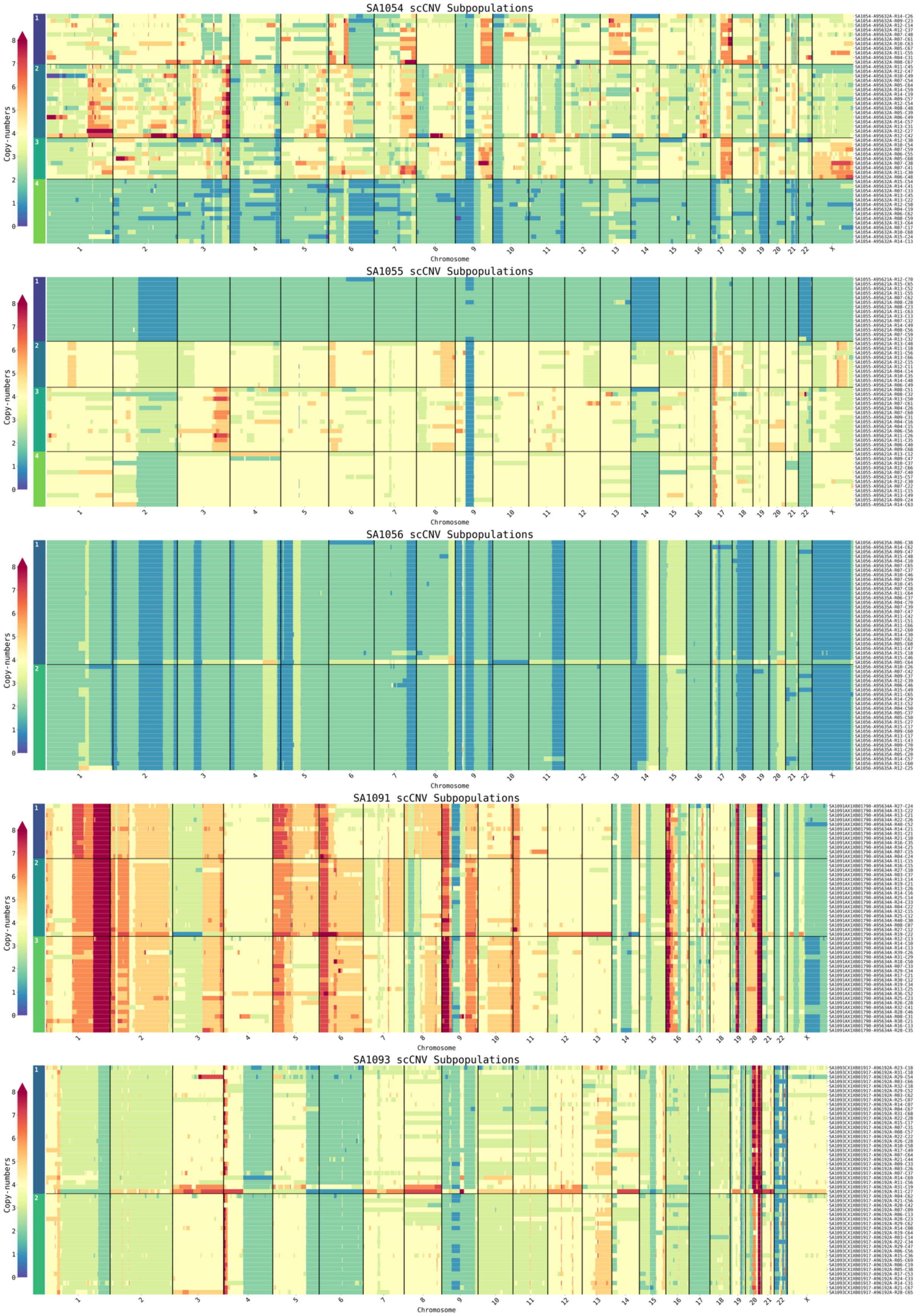
SA1054 scCNV Subpopulations
SA1055 scCNV Subpopulations
SA1056 scCNV Subpopulations
SA1091 scCNV Subpopulations
SA1093 scCNV Subpopulations
Copy-numbers
Chromosome

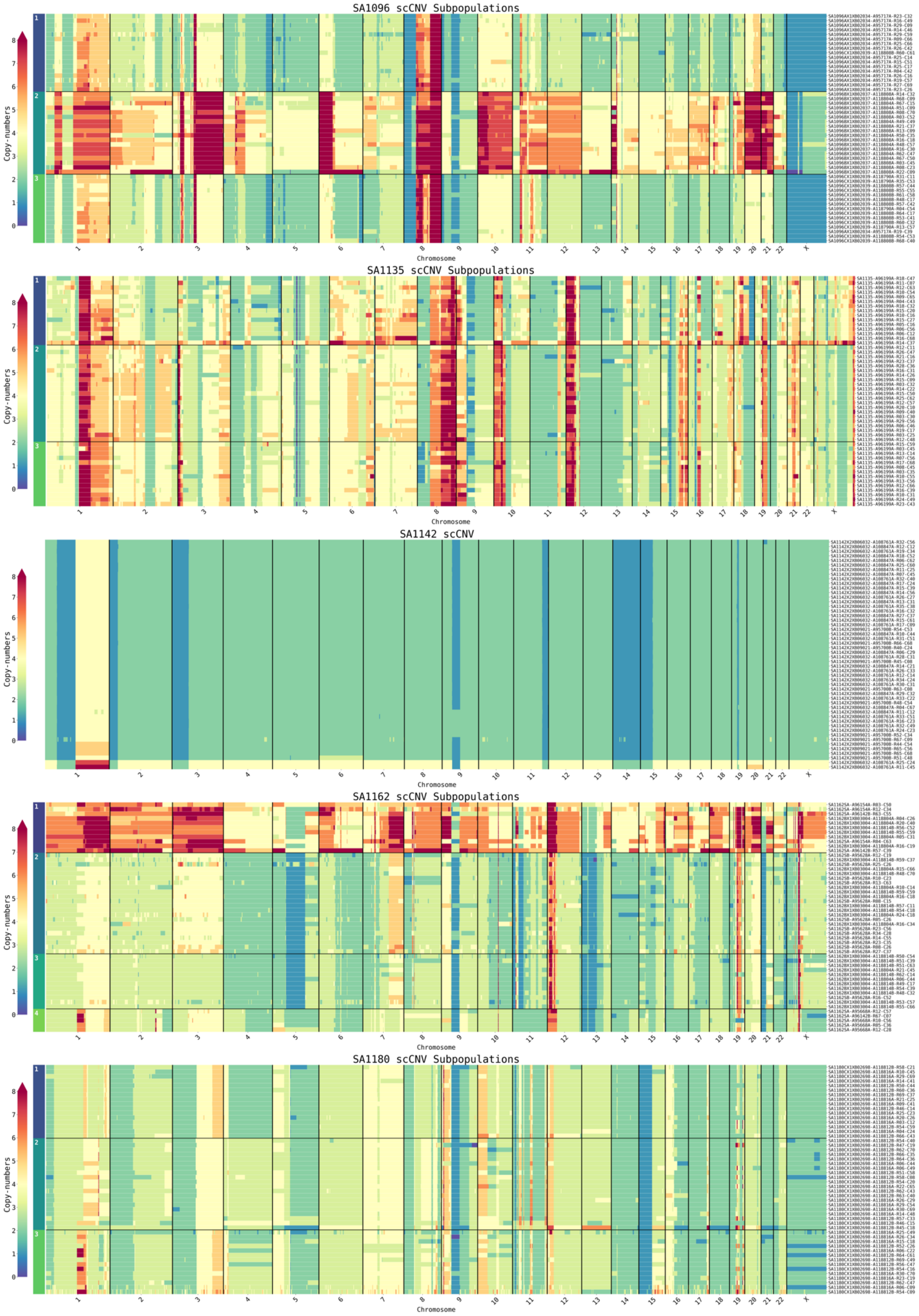

SA1096 scCNV Subpopulations
SA1135 scCNV Subpopulations
SA1142 scCNV
SA1162 scCNV Subpopulations
SA1180 scCNV Subpopulations
Copy-numbers
Chromosome

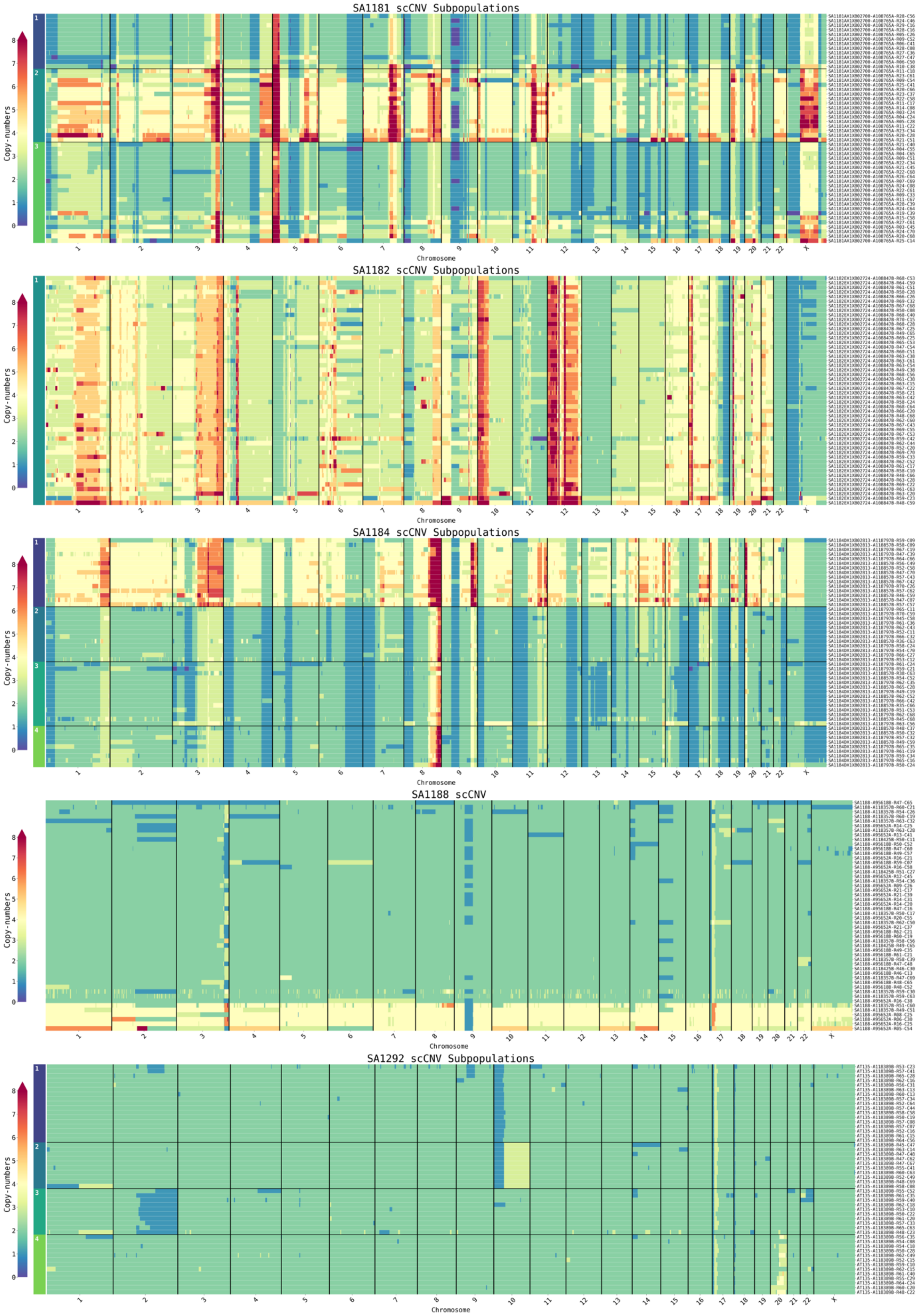


**Supplementary Figure S3.** Genome-wide scCNV profiles of a maximum of 50 randomly selected cells for Connolly2022 **(A)**, Du2021 **(B)**, Gnan2022 **(C)**, Laks2019 **(D)**, Massey2022 **(E)**,

Minussi2021 **(F)**, Takahashi2019 **(G)** and Funnell2022 **(H)** split by cell-phase (A,G) or subpopulation (B-F,H).
